# Individual structural features constrain the mouse functional connectome

**DOI:** 10.1101/613307

**Authors:** Francesca Melozzi, Eyal Bergmann, Julie A. Harris, Itamar Kahn, Viktor Jirsa, Christophe Bernard

**Author notes:** Corresponding authors: Christophe Bernard and Viktor Jirsa. Equal contribution first authors (Francesca Melozzi, Eyal Bergmann). Equal contribution last authors (Itamar Kahn, Viktor Jirsa, Christophe Bernard).

## Abstract

Whole brain dynamics intuitively depends upon the internal wiring of the brain; but to which extent the individual structural connectome constrains the corresponding functional connectome is unknown, even though its importance is uncontested. After acquiring structural data from individual mice, we virtualized their brain networks and simulated *in silico* functional MRI data. Theoretical results were validated against empirical awake functional MRI data obtained from the same mice. We demonstrate that individual structural connectomes predict the functional organization of individual brains. Using a virtual mouse brain derived from the Allen Mouse Brain Connectivity Atlas, we further show that the dominant predictors of individual structure-function relations are the asymmetry and the weights of the structural links. Model predictions were validated experimentally using tracer injections, identifying which missing connections (not measurable with diffusion MRI) are important for whole brain dynamics in the mouse. Individual variations thus define a specific structural fingerprint with direct impact upon the functional organization of individual brains, a key feature for personalized medicine.

**SIGNIFICANCE STATEMENT:** The structural connectome is a key determinant of brain function and dysfunction. The connectome-based model approach aims to understand the functional organization of the brain by modeling the brain as a dynamical system and then studying how the functional architecture rises from the underlying structural skeleton. Here, taking advantage of mice studies, we systematically investigated the informative content of different structural features in explaining the emergence of the functional ones. We demonstrate that individual variations define a specific structural fingerprint with a direct impact upon the functional organization of individual brains stressing the importance of using individualized models to understand brain function. We show how limitations of connectome reconstruction with the diffusion-MRI method restrict our comprehension of the structural-functional relation.

## INTRODUCTION

Structural connectivity (SC) refers to set of physical links between brain areas (Connectome, (1)) and constitutes an individual fingerprint in humans (2, 3). Since the connectome provides the physical substrate for information flow in the brain, it should impose strong constraints on whole brain dynamics. Functional connectivity (FC), in the context of resting-state functional MRI, refers to coherent slow spontaneous fluctuations in the blood oxygenation level-dependent (BOLD) signals measured in the passive awake individual. FC is commonly used to assess whole brain dynamics and function (4). Similar to SC, FC constitutes an individual functional fingerprint (5–7) and shows specific alterations during aging and in brain disorders (8). There is thus a strong correlation between the structural and the functional connectome. However, the causal relation between SC and FC remains unknown. Large scale brain modeling offers a way to explore causality between structural and functional connectivity. Combining experimental and theoretical approaches, we here unravel and quantify the degree to which the individual’s SC explains the same individual’s variations in FC.

We use The Virtual Brain (TVB), which allows building individual brain network models based on structural data (9). This brain network modeling approach operationalizes the functional consequences of structural network variations (10, 11) and allows to systematically investigate SC-FC relations in individual human brains (12– 15). If SC constrains FC, SC-based simulations of FC should match empirical FC within the bounds of validity of the metric. In primates and rodents, individual SCs are derived from diffusion MRI (dMRI). However, dMRI does not provide information on fiber directionality or synaptic details (distribution and type of neurotransmitter), and suffers from limitations, such as underestimation of fiber length and misidentification of crossing fiber tracks (16, 17). Given the imprecision of dMRI derived SC, it is difficult to estimate the validity of the simulations. This would require the knowledge of the ground truth connectome of an individual, which cannot be measured at present. However, the currently best gold standard can be derived in mice from cellular-level tracing of axonal projections (18), here named the Allen connectome. Although individuality is lost (the SC is a composite of many mice) and despite other limitations (19, 20), the Allen connectome provides details not available otherwise and in particular not available in humans. Focusing our attention on simulating mouse brain dynamics, we can thus use this detailed connectome to explore which missing features in the dMRI account for individual SC-FC relations. Specifically, we predict that fiber directionality and fine grain connectivity patterns should be key determinants.

Using dMRI data of 19 mice, we constructed 19 virtual mouse brain models (21) and compared predicted FC with empirical FC data acquired from the same mice during passive wakefulness (22). We found that individual SC predicts individual FC better than the dMRI-based averaged SC, and that predictions can be improved by considering fiber directionality, coupling weights and specific fiber tracks derived from the Allen connectome. We also found that hemispherical lateralization in the mouse connectome influences whole brain dynamics.

## RESULTS

We collected both dMRI and awake resting-state fMRI data (7 sessions per animal) from 19 hybrid B6/129P mice. We extracted SC from dMRI data to build individual virtual brains, which were imported into The Virtual Mouse Brain (TVMB), the extension of the open source neuroinformatic platform TVB (9) designed for accommodating large-scale simulations and analysis in the mouse, to generate *in silico* BOLD activity (21) using the reduced Wong Wang model (14, 23). We then compared simulated and empirical FC for each mouse in order to assess the power that an individual SC has to predict individual empirical FC derived from resting-state fMRI data (Figure S1). Further, SC was also obtained from the Allen connectome (our gold standard) in TVMB (21) to determine the contribution of information not available in dMRI-based SC. Experimental and simulated resting-state activity was characterized by a dynamical switching between stable functional configurations as revealed by the typical checkerboard patterns of Functional Connectivity Dynamics (FCD, Figure S2a and S2b), as observed previously (14, 24, 25). As expected, FCD varied across recording sessions (Figure S2b). In contrast, static Functional Connectivity (FC) was stable between experimental recording sessions (Figure 1A and Figure S2c). To compare the goodness of *in silico* resting-state dynamics against *in vivo* data, we needed a metric stable across experimental recording sessions in individual subjects, and thus we used the static FC for evaluating the Predictive Power (PP) of a SC, instead of FCD.

**Figure 1.**
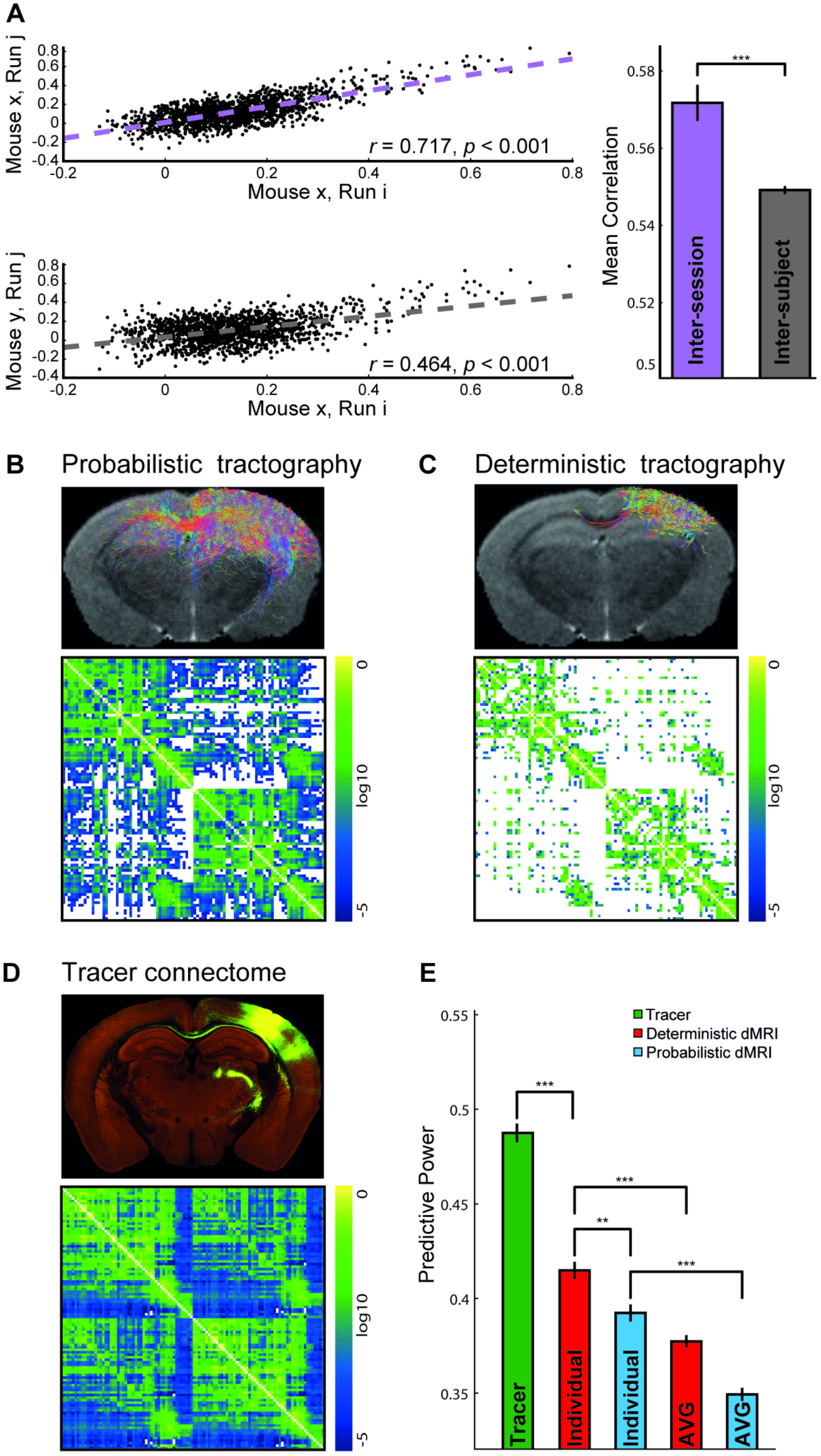
FC reliability was evaluated by comparing functional connectivity estimates between sessions. Representative scatter plots (*left*) show the correlations between sessions of the same mouse (*inter-session, top*) or between sessions of different mice (inter-subject, *bottom*). Quantification of inter-session and inter-subject similarities in the whole dataset (right) revealed that inter-session similarity is significantly higher than inter-subject similarity. Welch’s test ***P<0.001. (B) Probabilistic and (C) deterministic connections for the right barrel-related primary somatosensory cortex (SSp-bfd, *top*) and for the whole brain (*bottom*) of an individual mouse. (D) Tracer-based connections from SSp-bfd (*top*) and group tracer-based SC matrix (the Allen SC, *bottom*). (E) Predictive power of simulations using different types of tracer- and dMRI-based SCs. dMRI-based simulations were performed using individual or group-averaged dMRI (AVG). Welch’s test, Bonferroni corrected, **P<0.01 ***P<0.001. As in the following figures, the boxes, in panel A and E, extend from the lower (0.25, Q1) to the upper (0.75, Q3) quartile values of the data, with a line at the median. The upper whisker of each box extends to last datum less than Q3 + 1.5*(Q3-Q1); the lower whisker extends to the first datum greater than Q1 - 1.5*(Q3-Q1).

We first defined the upper bound of the PP. The correlation value calculated between any pair of empirical FC for each mouse provides us with an upper boundary of the PP, taking into account inter-session variability and other sources of noise that preclude 100% PP accuracy (7, 26). In keeping with human data (6, 27), we found a high inter-session correlation for each of the 19 mice, demonstrating stability across different recording sessions in a given mouse (Figure 1A). Inter-session correlations within the same animal were greater than inter-subject correlations, indicating that there is an individual functional organization per mouse, which may act as a functional fingerprint. Next, we sought to examine the extent to which individual functional connectomes correspond to individual structural connectomes.

### SC obtained with a deterministic algorithm is a better predictor of FC

Here we considered probabilistic (Figure 1B) and deterministic (Figure 1C) dMRI-based SCs, using iFOD2 (28) and SD_Stream (29) within Mrtrix3 software (29) tractography algorithms, respectively. SC obtained with the deterministic algorithm yielded a greater PP than the SC obtained with the probabilistic one (*PP*_*Individual–det*_ = 0.415 ± 0.005, *PP*_*Individual–prob*_ = 0.392 ± 0.005, 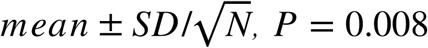 for the Welch’s test Bonferroni corrected, Cohen’s d=0.45, 95% CI=[0.19, 0.71], N=120; Figure 1E). The significative density difference in the two kinds of connectomes (*Density*_*Individual*−*prob*_ = 69 ± 1*%*, *Density*_*Individual*−*det*_ = 28.2 ± 0.2*%*, P = 3e^-20^ for the Welch’s test, Cohen’s d=12, 95% CI=[8, 17]), by itself, is not enough to explain the observed discrepancy in the PP. Connection density does not fully account for the predictive power of a connectome, but instead the relation depends on the connectome derivation (Figure S3). We argue that the observed difference in PP between deterministic and probabilistic processed connectomes depends on the proportion of false negative (FN) and false positive (FP) connections introduced by the two different algorithms. Zalesky and colleagues (2016) (30) show that the typical brain small-world topology is biased by the introduction of FP connections two times more than by the introduction of FN connections. In line with this finding, we attribute the difference in PP of the two connectomes to the detrimental role of FP connections, which are more likely introduced by probabilistic than deterministic tractography. However, deterministic tractography more likely overlooks some connections, introducing FN. This highlights the importance of preserving SC specificity (FN versus FP) versus SC sensitivity (FP versus FN) in the context of large-scale models. Namely, to preserve the global topology, specificity is more important as sensitivity in SC reconstruction. In the following, we compared deterministic SC-based simulated and empirical FCs.

### Individual SC is the best predictor of individual FC

Next, we found that individual SCs had a greater predictive power than the averaged SC (*PP*_*Individual*−*det*_ = 0.415 ± 0.005, *PP*_*AVG*−*det*_ = 0.377 ± 0.003, P = 3e^-9^ for Welch test Bonferroni corrected, Cohen’s d=0.86, 95% CI=[0.60, 1.12], *PP*_*Individual*−*prob*_ = 0.392 ± 0.005, *PP*_*AVG*−*prob*_ = 0.349 ± 0.004, P = 2e^-11^ for the Welch’s test Bonferroni corrected, Cohen’s d=0.97, 95% CI=[0.71, 1.23], N=120; Figure 1E), showing the importance of individual SCs. Although the Allen SC was obtained from hundreds of different mice, we found that it had a greater PP than individual dMRI-based SCs (*PP*_*Individual*−*det*_ = 0.415 ± 0.005, *PP*_*Tracer*_ = 0.488 ± 0.005, P = 4e^-21^ for the Welch’s test Bonferroni corrected, Cohen’s d=1.39, 95% CI=[1.05, 1.72], N=120; Figure 1D,E), suggesting that the tracer-based connectome includes structural information that is not present in dMRI, but which is central to explain the emergence of the functional connectome, even at the individual level. As the Allen SC was built from C57BL/6 mice, we verified the generality of our results in this strain (Figure S4a). Global signal regression, which improves structure-function relations and averaging recording sessions within each mouse (31), which reduces noise, increased the PP but did not alter the results (Figure S4b-c). Finally, splitting the recording sessions of each mouse, and submitting the data to a test-retest analysis revealed a close agreement between datasets (Figure S4d). Thus, our conclusions are strain- and preprocessing-independent, and reproducible.

### Importance of long-range connections and directionality

To identify the source of the systematic superior performance of the Allen SC, we focused on the major limitations of dMRI: (i) difficulty to resolve long axonal tracts, (ii) lack of information on fiber directionality and synaptic transmission and (iii) imprecise estimation of connection weights caused also by the impossibility to detect synaptic connections. Although synaptic properties are not available with precision at present, other parameters can be estimated. We tested the contribution of fiber length by filtering the Allen SC to include only fibers present in the dMRI-based SC (Figure 2A). We characterized the role of fiber directionality by symmetrizing the Allen SC (Figure 2A), asymmetrizing the dMRI-based SC (Figure 2B), and quantifying the impact of each manipulation (Figure 2C).

**Figure 2.**
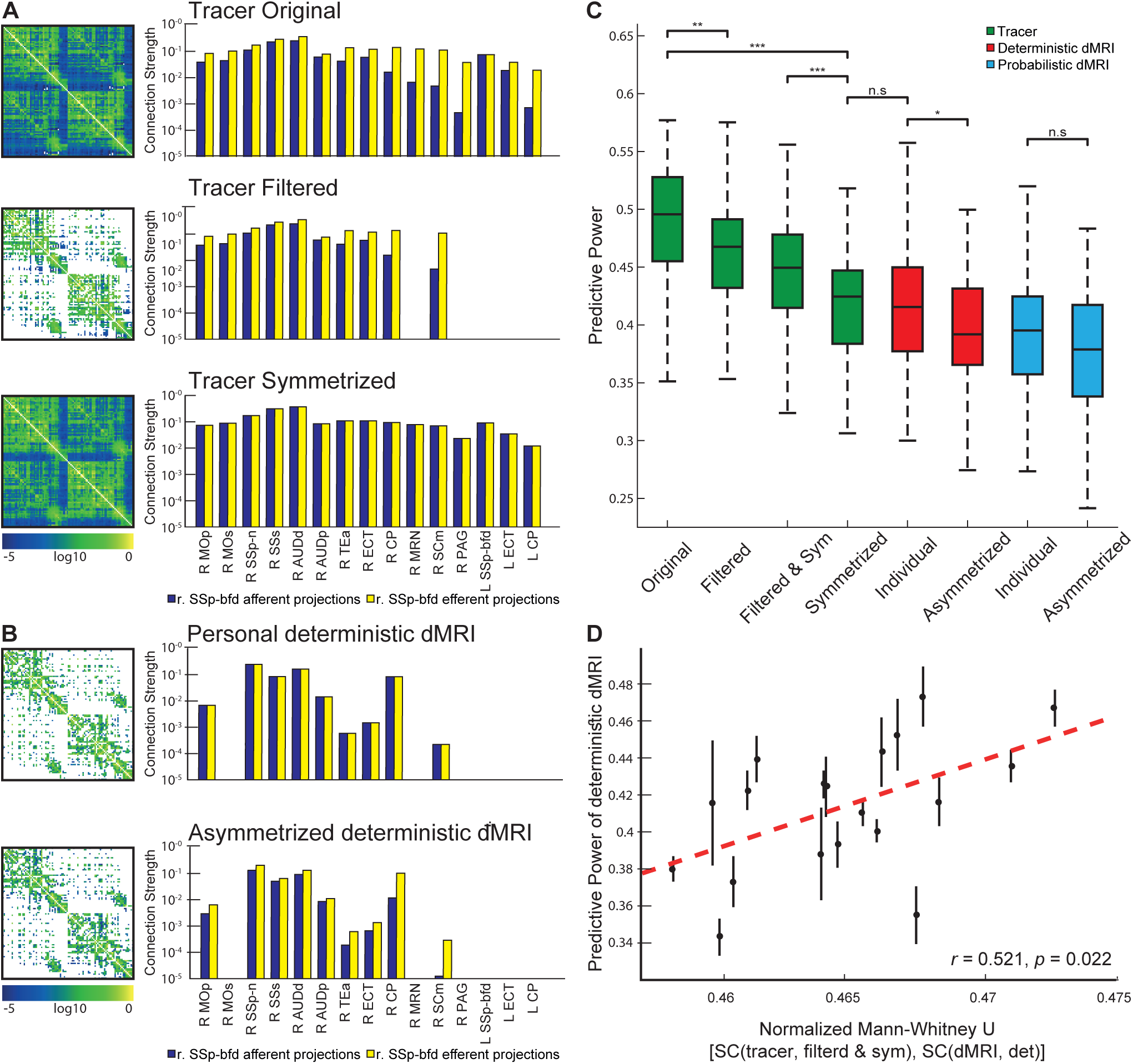
SC surrogates generated from the tracer-based SC (*left*) and representative connections of the SSp-bfd (*right*). The original, filtered and symmetrized SCs are shown in the top, middle and bottom rows, respectively. (B) A representative individual deterministic-dMRI SC (*left*) and SSp-bfd’s connections (*right*) before (*top*) and after (*bottom*) a-symmetrization. (C) Comparison between the performances of different surrogate SC demonstrates the effects of filtering and/or symmetrization of the Allen SC and a-symmetrization of the individual deterministic and probabilistic dMRI-based SCs. Welch’s test, Bonferroni corrected, *P<0.05, **P<0.01, ***P<0.001. (D) The relations between PP of filtered and symmetrized tracer SC and individual deterministic dMRI SCs are quantified through the normalized Mann-Whitney U-static and demonstrate significant correlation. The greater U-values represent greater similarity between individual deterministic dMRI SCs and the Allen SC.

Since dMRI fiber reconstruction reliability is inversely proportional to fiber length (16, 32, 33), dMRI SCs are sparser than the Allen SC (Figure 1B-C-D, S3a). To test the influence of the missing fibers in predicting FC, we built a filtered Allen SC (Figure 2A), which includes only the connections contained in at least one of the 19 deterministic dMRI SCs. The filtered connectome contains the 32% of the connections of the original tracer connectome, that are those captured by the dMRI-based deterministic processed connectomes. The connections that remain after the filtering operation are mainly those characterized by short-range length (Figure S3B): the averaged path length of the connections in the original and filtered tracer-based connectome is 5.40±0.02 mm and 3.57±0.03 mm, respectively. Figure 2C shows that the PP of the filtered Allen SC is lowerthan the original Allen SC (*PP*_*Tracerfiltered*_ = 0.461 ± 0.005, *PP*_*Tracer*_ = 0.488 ± 0.005, *P* = 0.002 for the Welch’s test Bonferroni corrected, Cohen’s d=0.49, 95% CI=[0.23, 0.76], N=120; Figure 2C), however it remains statistically greater than the PP of individual SCs (*PP*_*Individual*−*det*_ = 0.415 ± 0.005, P = 2e^-10^ for the Welch’s test Bonferroni corrected, Cohen’s d=0.92, 95% CI=[0.62, 1.20], N=120; Figure 2C). Thus, although connections overlooked by the dMRI method, which are mainly long-range connections, are important to explain FC, other important structural features present in the Allen SC are necessary to explain the discrepancy in PP between the tracer-based and dMRI-based connectomes.

We next focused on fiber directionality, since imposing bidirectional communication between regions connected with unidirectional links *in vivo* may affect FC. We used an approach based on surrogate SCs to test the role of directionality. Since the Allen SC contains directionality between regions, we removed this information by symmetrizing it (Figure 2A). Figure 2C shows that symmetrizing the Allen SC reduces its PP significantly (*PP*_*Tracersym*_ = 0.418 ± 0.004, *PP*_*Tracer*_ = 0.488 ± 0.005, P = 2e^-20^ for the Welch’s test Bonferroni corrected, Cohen’s d=1.36, 95% CI=[1.02, 1.68], N=120; Figure 2C), making it comparable to the PP of the dMRI-based SCs (*P =* 1.0 for the Welch’s test Bonferroni corrected, Cohen’s d=0.06, 95% CI=[-0.19, 0.32], N=120). This demonstrates that directionality is a key determinant of FC. It is notable that symmetrizing the filtered Allen SC led to a more modest reduction of the PP than the symmetrization of the original Allen SC (*PP*_*Tracersym*_ = 0.418 ± 0.004, *PP*_*Tracerfilteredsym*_ = 0.446 ± 0.004, P = 4e^-5^ for the Welch’s test Bonferroni corrected, Cohen’s d=0.62, 95% CI=[0.36, 0.87], N=120; Figure 2C). We argue that the PP difference can be explained by considering the amount of false positive introduced in the surrogate connectomes by the transformation: the filtering operation inserts FN connections, while the symmetrization operation inserts both FN and FP connections (34). It follows that the symmetrized and filtered connectome contains less FP than just the symmetrized connectome. Thus, as previously discussed for the tractography processing, introducing FP connections, as produced by the symmetrisation but not by the filtering, is more detrimental than the introduction of FN connections. To summarize when the tracer-based connectome is manipulated in order to remove the information not detected by dMRI, which is the inability to detect (i) the directionality of brain connections, as well as, (ii) some brain connections, especially the long-range ones, we found that the removal of the directionality information biases the predictive power of the connectome more than the removal of the connections not detected by the dMRI method.

We then took the complementary approach: enriching the dMRI-based SC with information on fiber directionality, i.e. asymmetrizing it. The results show that asymmetrizing the dMRI SCs does not increase, but rather decreases the PP (*PP*_*Individual*−*det*_ = 0.415 ± 0.005, *PP*_*Individual*−*det*−*asym*_ = 0.394 ± 0.005, *P* = 0.02 for the Welch’s test Bonferroni corrected, Cohen’s d=0.42, 95% CI=[0.16, 0.67], N=120; *PP*_*Individual*−*prob*_ = 0.392 ± 0.005, *PP*_*Individual*−*prob*−*asym*_ = 0.377 ± 0.005, *P* = 0.3 for the Welch’s test Bonferroni corrected, Cohen’s d=0.29, 95% CI=[0.04, 0.55], N=120; Figure 2B,C). We argue that the asymmetrization of the dMRI connectomes biased the PP because asymmetrizing a matrix is an ill-posed problem, since there is no unique solution (more details can be found in the Methods). In addition, there is no 1:1 correspondence between the connection strengths obtained with dMRI (axonal bundles) and Allen ones (axonal branches) since axons tend to branch more or less profusely when reaching their target zone, a feature that cannot be detected by dMRI.

### Connection strengths as key determinants of FC

The symmetric filtered Allen SC and the deterministic dMRI SCs have a similar structure: both matrices are symmetric and contain the same number of elements. Since the PP of the symmetric filtered Allen SC is still greater than the dMRI one, the difference can only result from dissimilarities in the values of the matrices’ entries, i.e. the connection strength values. Figure 2D shows that there is a significant relation between the normalized U-statistics of the Mann-Whitney test calculated between the filtered symmetric Allen SC and the individual dMRI SC and the PP of the latter (r=0.52, P=0.02). Namely, the more the distribution of connection strengths of the deterministic dMRI is similar to that of the Allen SC, the more reliable the predictions are.

### Specific refinement of individual dMRI connectomes

Since some afferent and efferent connections of specific areas may not be reliably reconstructed with dMRI, we examined whether refining dMRI SCs with more precise patterns derived from the Allen SC would improve the PP. For each deterministic dMRI SC, we substituted the non-zero incoming and outgoing connections of a specific region with the corresponding Allen SC projections, thus building a *hybrid* connectome (Figure 3A, S5).

**Figure 3.**
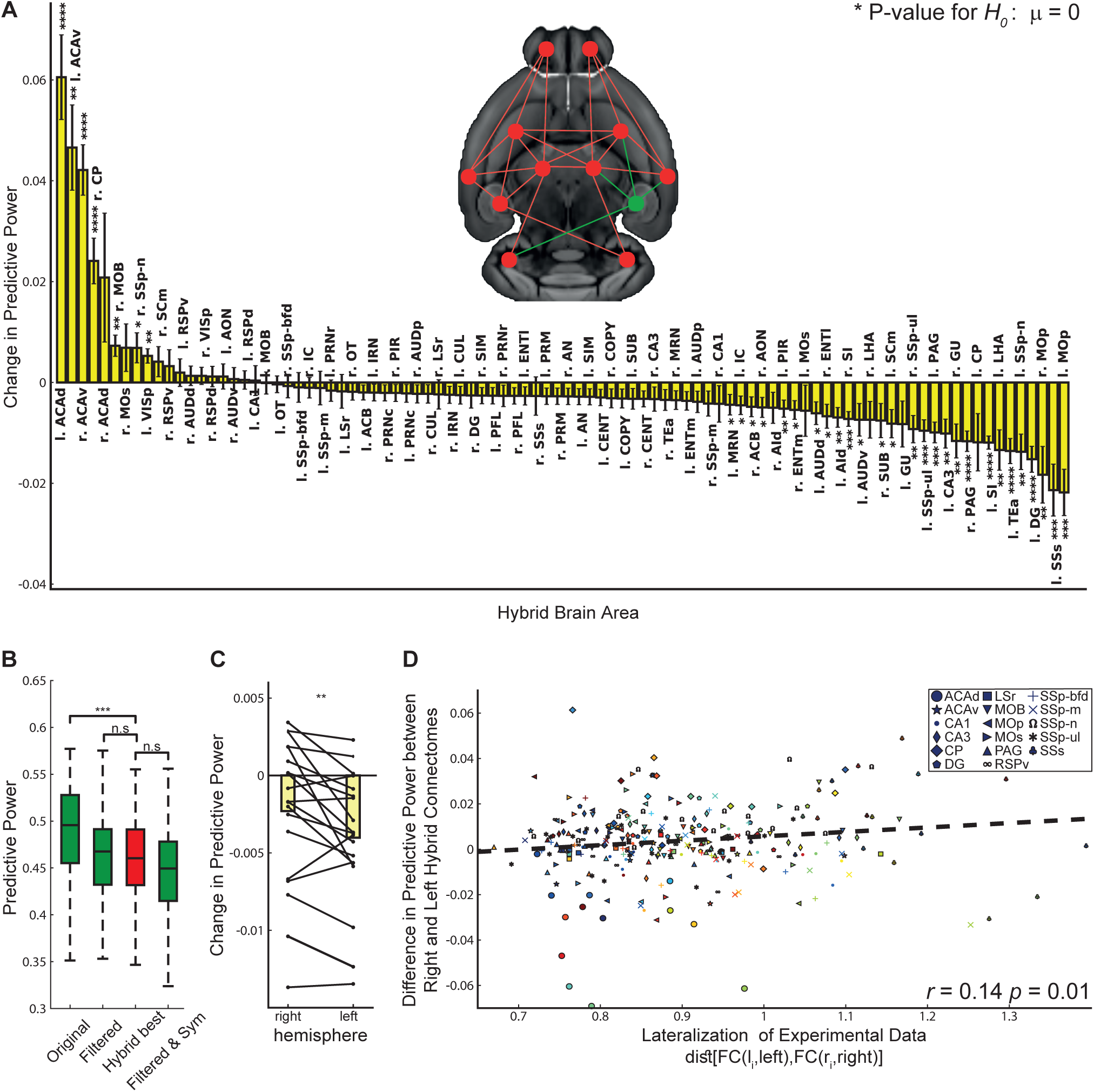
(A) An illustration of a hybrid SC. The connections between all areas are extracted from dMRI data (*red* nodes and links) except for the connections of a single area, which are obtained from the Allen SC (*green*). (B) The graph shows the difference in PP between hybrid and individual deterministic dMRI SCs. The change in PP is calculated as the difference between the PP of the hybrid SC and the individual dMRI SC averaged across all sessions and animals. P-values refer to the t-test for the null hypothesis that substituting the tracer connections of a given brain region in the dMRI connectome does not change the PP of the connectome. Nomenclature and abbreviations are listed in Table S1. (C) A Comparison between the PP of tracer-based and hybrid SCs revealed that connectomes (Hybrid Best), which were generated by replacing the connection of a single area in each mouse, predict experimental FC better than the filtered and symmetrized connectomes (Filtered & Sym). Welch’s test, Bonferroni corrected, **P<0.01. (D) Comparison between the change in PP following hybridization of left and right brain areas reveals that replacement of left areas’ connections decreased the PP more than right areas’ connections (paired t-test). (E) The Differences in PP between right and left hybrid SCs were correlated with the lateralization of experimental FC, which was quantified as the Euclidean distance between left and right functional connections. The closer the lateralization index is to 1, the more similar are the intra-hemispheric left and right area’s connections. Different colors label different mice.

When considering all mice, we found that substituting the anterior cingulate areas and the right caudoputamen connectivity patterns with the Allen SC projections significantly improved the PP of the connectome (left ACAd, improvement =0.047±0.006, t=7.23, P = 7e^-6^ for the T-test; left ACAv, improvement 0.032±0.006, t=4.96, *P* = 0.002 for the T-test; right ACAv, improvement=0.028±0.003, t=7.58, P = 1e^-4^ for the T-test; right CP, improvement=0.018±0.003, t=6.42, P = 5e^-6^ for the T-test; Figure 3B), suggesting that both regions are poorly resolved by dMRI in mice. Importantly, the majority of substitutions decreased the PP (Figure 3B).

For each individual SC, we identified the region in which replacement of its dMRI-connections with the Allen ones generates a new connectome, hybrid^best^, which has the best PP improvement as compared to the other hybrid connectomes (Figure S5). Figure 3C shows that the PP achieved by hybrid^best^ is statistically indistinguishable to the one achieved by the filtered Allen SC (*P* = 1.0 for the Welch’s test Bonferroni corrected, Cohen’s d=0.008, 95% CI=[-0.25, 0.25], N=120). In other words, it is sufficient to replace in the dMRI SC the connections of one particular region with the corresponding Allen ones, to get a similar prediction, which is specific for each mouse.

### The asymmetric mouse brain

Finally, we sought to estimate the potential contribution of asymmetric transhemispheric connectivity. Figure 3D shows that there is a considerable improvement in the PP of hybrid SCs when using connections from the right hemisphere, as compared to those from the left one. The Allen connections have been estimated using unilateral injection in the right hemisphere (18). Since no tracer injections were done in the left hemisphere, TVMB uses a mirror image of the right hemisphere to build the left one (21). This suggests that the tracer-based intra-hemispheric connectivity predicts better right intra-hemispheric functional behavior than the left one, as demonstrated in Figure S6a. Figure 3E shows that there is a significant relation between hemispheric lateralization in the functional connectomes and the improvement in PP when the right and left homotopic tracer area’s connections are introduced in the dMRI SC (*r* = 0.14, *P* = 0.01). Namely, the more functional connections are asymmetric, the more the PP decreases when using the right hemisphere connections to build the left ones. These results suggest that connectivity asymmetry impacts brain dynamics and that it is region- and mouse-specific.

### Hemispherical lateralization of the mouse brain

Figure 3E shows that the region demonstrating the greatest lateralization in terms of functional connectivity in individual mice is the supplemental somatosensory area (SSs). Figure 3B shows that when we introduce the mirror image of the right SSs into the dMRI SC, the predictive power is considerably decreased, which means that the mirror image of the right SSs poorly represents the true left SSs. We thus focused on the SSs area. If SC drives FC, we predicted that introducing in the tracer-based connectome the detailed left SSs connections, instead of using the mirror image of the right SSs ones, would increase the PP of the connectome. We first performed tracer injections in the left SSs and determined the projection pattern. As predicted, we found evidence of an asymmetric distribution of fibers between the left and right SSs (Figure 4A). To test whether these structural differences were sufficient to explain the functional ones, we introduced the connections of the left SSs into the tracer connectome and obtained a statistically greater PP as compared to the ones of purely mirrored connectomes built from the injection experiments performed in the right SSs (Figure 4B). Next, we introduced the left connections of the SSs into the dMRI-based SCs (hybrid connectome), and, as predicted, we found a greater PP as compared to using the mirror image of the right connections of the SSs as shown in Figure 4C (between the 14 experiments performed in the right SSs we consider the one whose injection location is more similar to those used in the left SSs injection experiment). Finally, since our previous results demonstrate that the lateralization is animal-dependent, we sought to examine whether lateralized FC is supported by lateralized SC and found that the improvement of the PP following hybridization of left SSs dMRI connections is indeed proportional to the degree of functional lateralization (Figure 4D). Together, these results show that the mouse brain is structurally lateralized, and that this lateralization impacts whole brain dynamics at the individual subject-level.

**Figure 4.**
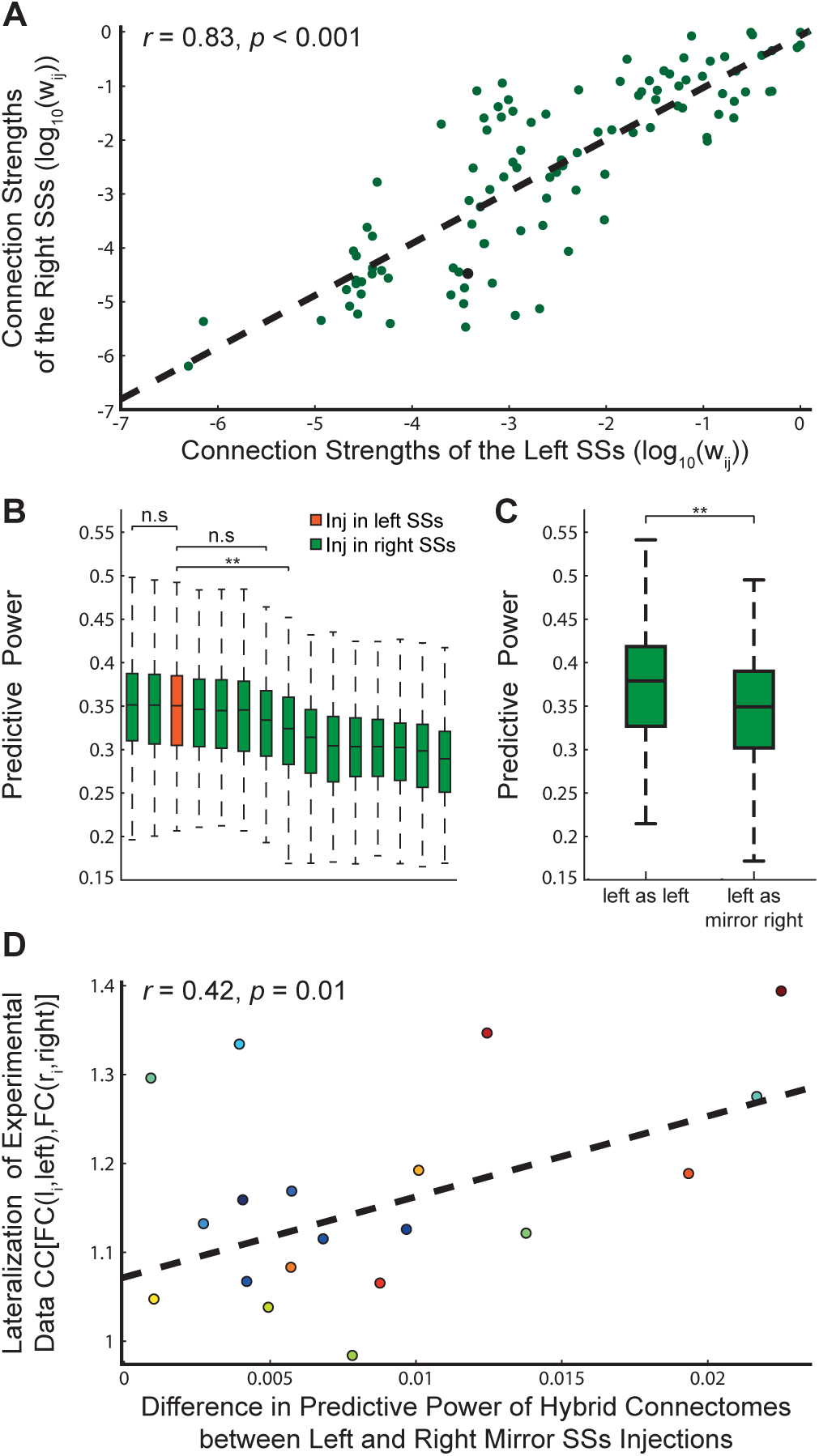
(A) Relation between fiber projections of the left and right SSs. (B) The height of the bars represents the predictive power of tracer-based connectomes built using just one injection experiment per area. This procedure differs from the general tracer building procedure used in this work since generally the connections of each region are calculated as the average of the results of all injection experiments performed in that region. The height of the orange bar is the predictive power of the tracer-based connectome whose left SSs connection are reconstructed from the tracer injection experiment in the left SSs. The green bars are the predictive power of the tracer-based connectomes whose left SSs connections are built as the mirror image of the connections measured from each of the 14 experiments performed in the right SSs area. The statistical difference between the bars is assessed through the Welch’s test, Bonferroni corrected, **P<0.01. (C) Comparison between the predictive power of hybrid connectomes built as a dMRI-based connectome with left SSs tracer connections reconstructed from the tracer injection experiment in the left SSs or as the mirror image of right tracer connections retrieved from the right SSs experiment whose injection coordinated are closer to those of the left SSs injection experiment. (D) The Differences in PP between true left and mirror right SSs hybrid SCs are correlated with the lateralization of experimental FC.

## DISCUSSION

Our results provide direct evidence of a type of causality between SC and FC, in the sense that individual structural connectomes predict their functional counterparts better than the dMRI-based averaged connectomes. Previous studies utilized the Allen Mouse Connectivity Atlas to study structure-function relations at the group level using voltage-sensitive dyes (35) and FC (6, 22, 36). In addition, a recent work in rats (37) used TVB to simulate FC based on SC and found strong correlation at the group level. A similar finding has been reported in humans (38). Here we compared structure-function relations in individual mouse brains and we used the detailed Allen connectome as a gold standard to identify regions and connections that play a preeminent role in the emergence of individual brain dynamics. We showed that, similar to humans (6), intra-mice FCs are more stable than inter-mice FCs (Figure 1A). We propose that the emergence of individual features in the functional data is, at least partially, driven by the individual’s structural connectivity with stable features encoded in the connectome (Figure 1E). In addition to connectome specificity, other structural data features may drive functional inter-subject variability including foremost regional variance such as synaptic receptor type and density (39), but also methodological variations such as parcellation differences (3). Notwithstanding, we cannot exclude that the variations in hemodynamic response functions (HRF) across animals and brain location affect SC-FC relations, as it has been shown to contribute to individual variability in human FC estimation (40). In this study we aimed at reducing this variability by scanning awake mice, reducing the confounding effects of anesthesia and allowing collection of data over multiple sessions per mouse (41, 42). Moreover, we used spin-echo echo planar imaging (EPI), which is more sensitive to microvasculature relative to gradient-echo EPI, especially at high magnetic fields (43), further reducing the variability of HRF (40). Finally, HRF variability can increase the number of false positives in FC in empirical data, but it cannot explain differences in predictive power of simulated data obtained from different structural connectomes.

Virtualizing different structural connectomes, we found that group tracer-based connectomes predict empirical FC better than individual dMRI-based connectomes, and while we argue that this difference can be explained by better measurements of long-range connections, fiber directionality and connection weights, we cannot rule out that it is caused, at least partly, by the compromised DTI data quality of *in-vivo* measurements. As compared to *ex-vivo* studies (44), we designed our DTI sequences with relatively low number of gradient directions, anisotropic voxels and relatively low resolution. Together with motion-related noise, these factors reduce the quality of tractography (45), and may contribute to the lower performance of dMRI-based SC in predicting empirical FC. Nevertheless, the DTI sequence was designed this way to allow in-vivo measurement in mice which is comparable to human data which will support future mechanistic investigations of SC-FC relations. The dMRI-connectome reconstruction could be improved by using more sophisticated anatomical constraints in the tractography pipeline (e.g. ACT method, (46)) in addition to the basic segmentation in regions of interest.

The detrimental role of false positive (FP) connections in the connectome topology has been previously explored by (30) and (34) analyzing, respectively, the effect of FP as introduced by probabilistic tractography and overlooking connections’ directionality. In line with these findings, we showed that the introduction of FP connections biases the connectome predictions. We found the dMRI-based connectomes processed with the deterministic tractography have a statistically greater PP than those processed with probabilistic algorithms. Since the observed difference in PP is not directly related with the difference in connections density (Figure S3), we argue that the difference in PP is driven by the different characteristics of the connections overlooked by both types of tractography processing: more FP and less FN in the case of probabilistic processed connectivity, and conversely in the case of deterministic processed connectivity. This highlights that brain dynamics predictions are more accurate if connectome specificity is preserved, even at the cost of sensitivity, as it is the case of deterministically processed connectomes.

When processing the tracer-based data, the probabilistic computational model used to construct the original Allen connectome (18) may introduce several false negative connections, resulting in a low connection density reconstruction (35-73%), whilst others reported a 97% density (19, 20). Here, we have used the Allen connectome builder interface, which implements a deterministic approach to reconstruct whole brain connectivity (21), leading to a 98% density of connections. Still, as shown in Figure 2B, the introduction of FN connections (filtered tracer-based connectome) does not dramatically influence the PP of the connectome.

The main drawback of the Allen connectome is that it has been obtained from hundreds of different mice, thus blurring individual variability. We found that replacing most individual dMRI connections with Allen connections reduces the PP. However, in some regions such as the anterior cingulate and the caudoputamen, group-level Allen connections outperform individual dMRI connections. This finding can be explained by the fact that connections from the anterior cingulate are difficult to resolve as this area is located in the midline brain region, where the cortex folds, resulting in an abrupt change in fiber directionality. Moreover, the axons make sharp turns around the corpus callosum while the extraction algorithm assumes a systematic continuation of the vector direction. The connections of the striatum are often short and, due to its multipolar organization without a clear gradient orientation limiting fiber reconstruction. These considerations apply to mouse strain used here. They cannot be translated as such to other species. But the conceptual framework we introduce shows how to take into account detailed connectivity patterns (if available) to improve the analysis of the SC-FC relationship. In humans, the release of full connectomes obtained postmortem at ultrahigh resolutions (47) constitutes an important step in this direction. Detailed connectivity patterns from non-human primates may also be used to build a high resolution “human” SC (48, 49).

Although the Allen connectome was obtained from C57BL/6 mice, brain dynamics of hybrid F1 mice could be predicted by the Allen connectome, suggesting that the structural organization of the mouse brain was not impacted by out-breeding. Findings from hybrid mice are considered more generalizable to other strains (50), thus suggesting that the pattern observed here is not strain-specific. Nonetheless, since the genetic background affects the behavioral phenotype (51), it will be important to systematically assess these findings in mouse strains where this aspect is directly manipulated.

The Allen SC includes directionality and long-range connections, which are not well (or at all) resolved by dMRI. However, the removal of the connections not resolved by dMRI-based connectomes, mostly those characterized by long-range length, is not sufficient to explain the discrepancy between the tracer-based and dMRI-based predictive power. In addition, we showed that removing the directionality information from the tracer-based connectome, that it is symmetrizing the connectome, thus introducing FP and FN connections, worsens the predictive power more than the filtering operation, that consist in introducing just FN connections (34). This shows the key role of connections directionality in predicting brain dynamics; and it confirms our results on tractography algorithm processing: FP connections biases the predictive power ability of the connectome more than FN. Finally, analyzing the connection strength differences between the dMRI and tracer-based connectome, we have showed that connection strengths are the main determinant of these dynamics, and consequently of individuality (Figure 2D).

An unexpected result was the important role played by the transhemispheric asymmetry of connections. This finding is consistent with calcium imaging studies reporting such asymmetry in rodents (52). By comparing injections between left and right hemispheres, we confirmed our prediction that the approximation of left area connections as compared to right area connections, necessary in the tracer-based connectome reconstruction, significantly affect the predictive power of the connectome. Moreover, we showed that the bias introduced by this approximation is proportional to the degree of the individual animal’s functional lateralization.

We quantified PP using the link-wise Pearson correlation across experimental and simulated FC. This choice of metrics has limitations linked to the stability of the functional data features during the time window considered. Other choices would have been possible, as for example comparing specific features of the FC (e.g. functional hubs, graph theoretical characteristics, subnetwork structures, etc.) or evaluating the dynamical evolution of the functional links, e.g. FCD or FMC (see Method section). Our definition of the PP, broadly accepted in literature (12–14, 37, 53–56), responds to the need to quantify the ability of a model to reproduce the functional brain behavior globally, and not its specific features. However, overlooking the informative content encoded in the dynamics of the FC is a limitation of our study. FMC is one means of quantifying the FCD globally via its dynamics of the functional links, but has proven to be too variable across resting state sessions within the same animal (Figure S2). This variability limits the possibility to compare simulated and experimental FMC, and to use it as a PP metric. Notwithstanding, we are omitting FCD features from the PP evaluation, these features are integral to the simulations (Figure S2).

Another limitation of our work is the assumption of region invariance, that is we use the same model, as well as the same parameters, to describe the activity of all the brain regions, both cortical and subcortical ones. The only symmetry breaking between the virtual brain areas is their integration in the network, i.e. their anatomical connections. It follows that in this approximation the connectome acquires a central role. This centrality allows us to trace back all the prediction differences obtained in the virtual mice to the connectome used, as it is our aim. However, we do acknowledge the need to introduce regional specificity and variance into the brain models. Future evolutions of large scale brain models should include such specificities.

Progress in connectomics enabled the development of large-scale brain models to study brain function in health and disease (12, 57). Although individual whole brain modelling has a potentially high translational value for the benefit of patients (15, 58, 59), the entire approach relies on the extent to which individual differences in structural connectomes determine the emergent network dynamics and consequent neuroimaging signals. Although SC does not provide enough information to predict an epileptogenic zone in humans (60), our work shows that using more precise information (e.g. obtained from tracer injections in non-human primates) to consider directionality, synaptic weights and poorly-resolved dMRI connections, will increase the predictive power. As for clinical applications the value of a generalized model is measured by its utility in individuals (15, 57), our results bear a significant promise in this domain as they demonstrate and quantify the predictive capacity of SC and FC variability.

In conclusion, we identified key individual structural features (fiber directionality, connection strength and patterns, and inter-hemispherical asymmetry), which are relevant to predict the emergence of the functional patterns during resting state in mice. Our results strongly suggest the existence of a causal relation between the structural and the functional connectome. Although the detailed structural results presented here are species-specific, our conceptual framework applies is species-invariant and can now be exploited in humans for individual diagnosis and clinical decision making.

## MATERIALS AND METHODS

Additional details on materials and methods are provided in the SI appendix.

### Structural and functional experimental data

19 male hybrid non-anesthesized mice were scanned using a 9.4 Tesla MRI to obtain structural information and resting state functional data, 6.31±0.82 (mean±SD) sessions of 15.7±4.4 minutes length (mean±SD) per mice. The structural data have been processed using using MRtrix3 software(29). To obtain the tract streamlines we integrated the field of orientation probability density using both deterministic (SD_Stream, (29)) and probabilistic (iFOD2, (28)) algorithms in order to obtain, respectively, deterministic and probabilistic processed dMRI-based connectome. Tracer-based connectome was built through the Allen Connectivity Builder pipeline (21), impelemented in The Virtual Brain software (9). The pipeline allows to manipulate the anterograde tracer experiments performed at the Allen Institute (18) to reconstruct a tracer-based mouse connectome and a related brain volume.

### Surrogate connectomes

In order to test different hypotheses about what could be the connectivity properties that give rise to the observed discrepancies in the simulated dynamics, we built different kinds of surrogate connectomes: a dMRI-based averaged connectome (averaging the 19 dMRI based connectomes), a filtered tracer-based connectome (filtering out the connections not detected in the dMRI-connectomes), a symmetrized tracer-based connectome, asymmetrized dMRI-based connectomes and hybrid connectomes (dMRI-based connectomes whose the connections of one area are replaced with the tracer-based connections of that area).

### Simulate resting state dynamics

We simulate resting state dynamics using the connectome-based model approach as implemented in The Virtual Brain software (9, 65). In particular, we used the reduced Wong Wang model (23, 13) in the bistable configuration in order to reproduce the dynamical switching of the functional connections (14). We transform the simulated synaptic activity in BOLD signal using the Balloon-Windekessel method (56, 57).

### Analysis

The emergence of the functional organization in the experimental and simulated brains was analyzed through the static Functional Connectivity (FC), the Functional Connectivity Dynamics (FCD) and the Functional Meta Connectivity.

We quantified the *Predictive Power (PP)* of a given connectome *c* as the Pearson correlation between the simulated FC, obtained using that connectome *c*, and the FC arranged during resting state experimental recordings.

In order to assess the significance of the difference in PP of differently derived connectomes we used the p-value calculated through the Welch’s test; we corrected the p-values for multiple comparisons using the Bonferroni correction. We measure the effect size using the Cohen’s d and we calculated the 95% confidence intervals (CIs) using the estimation stats framework as described in (67) and available at https://www.estimationstats.com/.

## Acknowledgements

Supported by ANR Connectome ANR-17-CE37-0001 (to F.M., V.J. and C.B.), European Union’s Horizon 2020 Framework Programme for Research and Innovation, Award ID: 785907 (HBP SGA2) to V.J., Israel Science Foundation (770/17; to I.K.), National Institutes of Health (1R01NS091037; to I.K.), Adelis Foundation (to I.K.) and Prince Center (to I.K.).

## SI APPENDIX

### 1. Animals and Surgical Procedures

All procedures were conducted in accordance with the ethical guidelines of the National Institutes of Health and were approved by the institutional animal care and use committee (IACUC) at Technion. 19 male first generation hybrid mice (B6129PF/J1, 9-12 weeks old) were implanted with MRI compatible head-posts using dental cement as previously described (22). After 3 days of recovery, the animals were acclimatized to extended head fixation. This training included 5 handling sessions performed over 3-5 days, and 4 daily acclimatization sessions inside the MRI scanner. In each acclimatization session, mice were briefly anesthetized with isoflurane (5%), and then head-fixed to a custom-made cradle for gradually longer periods (2, 5, 10, 25 min). Subsequently, mice underwent seven 45 min long awake imaging sessions, and one diffusion tensor imaging (DTI) session under continuous isoflurane anesthesia (0.5-1%). A second group that included 7 male inbred C57BL/6 mice (11-16 weeks old) was operated and scanned according to the same protocol.

Experiments involving mice were approved by the Institutional Animal Care and Use Committees of the Allen Institute for Brain Science in accordance with NIH guidelines. For left side injections into SSs, surgical procedures were followed as described in (18). In brief, a pan-neuronal AAV expressing EGFP (rAAV2/1.hSynapsin.EGFP.WPRE.bGH, Penn Vector Core, AV-1-PV1696, Addgene ID 105539) was used for injections into wildtype C57BL/6J mice at postnatal day 56 (stock no. 00064, The Jackson Laboratory). SSs was targeted using stereotaxic coordinates from Bregma (AP: −0.7, ML, −3.4 and −3.9) and from brain surface (DV: 1.66). rAAV was delivered by iontophoresis with current settings of 3 µA at 7 s ‘on’ and 7 s ‘off’ cycles for 5 min total, using glass pipettes (inner tip diameters of 10–20 µm). Mice were perfused transcardially and brains collected 3 weeks post-injection for imaging using serial two-photon tomography, using methods as previously described for the Allen Mouse Connectivity Atlas (18).

### 2. Data acquisition (fMRI and diffusion-MRI)

MRI scans were performed at 9.4 Tesla MRI (Bruker BioSpin GmbH, Ettlingen, Germany) using a quadrature 86 mm transmit-only coil and a 20 mm loop receive-only coil (Bruker). Mice were shortly anesthetized (5% isoflurane) before mounted on the cradle. After acquisition of a short low-resolution rapid acquisition process with a relaxation enhancement (RARE) T1-weighted structural volume (TR = 1500 ms, TE = 8.5 ms, RARE-factor = 4, FA = 180°, 30 coronal slices, 150 × 150 × 450 μm^3^ voxels, no interslice gap, FOV 19.2 × 19.2 mm^2^, matrix size of 128 × 128), four spin echo EPI (SE-EPI) runs measuring BOLD fluctuations were acquired (TR = 2500 ms, TE = 18.398 ms, 200 time points, FA = 90°, 30 coronal slices, 150 × 150 × 450 μm^3^ voxels, no interslice gap, FOV 14.4 × 9.6 mm^2^, matrix size of 128 × 128). In addition, mice underwent another session under anesthesia to acquire high resolution T2 image (TR = 6000 ms, TE = 8.8 ms, RARE-factor = 16, FA = 180°, 36 coronal slices, 100 × 100 × 400 μm^3^ voxels, FOV 16 × 16 mm^2^, matrix size of 160 × 160, 10 averages) and diffusion tensor imaging data (DTI) with a diffusion-weighted spin-echo echo-planar imaging (EPI) pulse sequence (TR = 9000 ms, TE = 21.68 ms, Δ/ δ=11/2.6 ms, 4 EPI segments, 30 gradient directions with a single b-value at 1000 s/mm^2^ and three images with b-value of 0 s/mm^2^ (B0), 36 slices, 100 × 100 × 400 μm^3^ voxels, FOV 16 × 16 mm^2^, matrix size of 160 × 160, 2 averages). Each DTI acquisition took 39.6 min.

### 3. Data processing

#### Intrinsic functional connectivity data

fMRI data preprocessing procedure was validated in a previous study (22). Briefly, the first two time points were removed for T1-equilibration effects, slice-dependent time shifts were compensated, head motion was corrected using rigid body correction, volumes were registered to a downsampled version of the Allen Mouse Brain Atlas, and data underwent intensity normalization. Then, motion scrubbing procedure was applied to remove motion-related artifacts as previously shown. Rigorous censoring criteria were used including frame displacement (FD) of 50 μm and temporal derivative root mean square variance over voxels (DVARS) of 105% of median. An augmented temporal mask of 1 frame before and 2 frames after detected motion was used and sequences of less than 5 included frames were also censored. Runs with less than 50 frames, and sessions with less than 125 frames (5.2 mins) were excluded. The average number of included sessions per mouse was 6.31±0.82 (mean±SD) for the F1 hybrid mice and 3.71±2.21 for the C57BL/6 inbred mice. Total included time per session was 15.7±4.4 (minutes per session, mean±SD) and 11.41±3.67, respectively.

After motion scrubbing, resting-state fMRI specific preprocessing procedure was applied including demeaning and detrending, nuisance regression of 6 motion axes, ventricular and white matter signals and their derivatives, temporal filter (0.009 < f < 0.08 Hz), and spatial smoothing (Gaussian kernel with FWHM of 450 μm.) The C57BL/6 group was preprocessed both with and without global signal regression to test the effects of this procedure on structure-function relations.

To estimate functional connectomes, we build a parceled volume with a resolution compatible with the fMRI technical constraints by manipulating the Allen Mouse Brain Connectivity Atlas (18) downloaded through The Virtual Brain (9, 21). The volume was registered to the space of the functional data (‘target.nii.gz’) using the nearest neighbor interpolation (FLIRT software, (61)). The parcellation was reduced only to the areas where the SNR was higher than 12, and that had a volume greater than 10 voxels (>0.1mm^3^). Finally, very anterior and posterior areas, such as the main olfactory bulb and cerebellum, were excluded from the parcellation due to registration problems and susceptibility artifacts associated with the head-post implantation. Once the parcellation volume was built, mean BOLD signals were extracted from the voxels composing each parcel, and correlations were calculated from included frames only (based on motion scrubbing).

#### Diffusion-MRI data

We processed diffusion-MRI data using MRtrix3 software(29).

The fiber orientation distribution of each voxel was estimated using the Constrained Spherical Deconvolution (CSD, (62)). To obtain the tract streamlines we integrated the field of orientation probability density using both deterministic (SD_Stream, (29)) and probabilistic (iFOD2, (28)) algorithms; in both cases, the tracts number was set to 100 million. The streamlines were then filtered using the SIFT algorithm (63) which selectively reduces the number of tracts exploiting the fiber orientation density information obtained through the CSD in the previous step. The filtered tracts of the right SSp-bfd obtained with probabilistic and deterministic algorithm, for an illustrative mouse, are shown in Figure 1B and 1C respectively. We defined seed regions using the Allen Mouse Brain Connectivity Atlas (18) obtained through the The Virtual Brain (9, 21); after registering the volume in the individual mouse diffusion space, we reduced the parcellation only to those areas whose volume was greater than 250 voxels (>1.125mm^3^). Using the deterministic and probabilistic streamlines and the node parcellation image, we generated a connectome. The connection strength between each pair of nodes was defined as the streamline count between the two nodes scaled by the inverse of the volumes of the two areas. A radial research was performed to assign each streamline end point to a given node. If no node was found inside a sphere of 1 mm radius, the streamline was not assigned to any node. We excluded all self-connections by setting the diagonal elements of the connectome to zero and normalized all connection strengths between 0 and 1. Then, we repeated this procedure for all mice. An example of personalized connectome obtained with probabilistic and deterministic algorithm is shown in Figure 1B and 1C, respectively.

#### Tracer data

The recent updates of The Virtual Brain software (9, 21) allows us to manipulate the anterograde tracer experiments performed at the Allen Institute (18) in order to obtain a very precise mouse connectome. Specifically, we define the link between two brain regions according to the anterograde tracing information provided by the Allen Institute of Brain Science and presented in the work of Oh et al., 2014 (18). In the latter, the axonal projections from a given region are mapped by injecting in adult male C57Bl/6J mice the recombinant adeno-associated virus, which expresses the EGFP anterograde tracer. The tracer migration signal is detected with a serial two-photon tomography system. This approach is repeated systematically in order to collect the information on the tracer migration from several injection sites in the right hemisphere to target regions in both ipsilateral and contralateral hemispheres; for each injection sites several experiments are run and distinct measures are accomplished. Through the Allen Connectivity Builder interface in TVB we define the connection strength between source region i and target region j as the ratio between the projection density (the number of detected pixels in the target region normalized on the total number of pixels belonging to that region) and the injection density of the source region (the number of infected pixels in the source region normalized on the total number of pixels belonging to that region). The TVB pipeline, once downloaded the raw data, i.e. projection density and injection density for experiment, from the Allen dataset, manipulate the data in order to build a connectome as described in Melozzi et al., 2017 (21).

Unless otherwise specified, the tracer-based connectome is built averaging experiments performed injecting the tracing compound in the areas in the right hemisphere.

One of the main differences between tracer and diffusion-MRI technique is the spatial resolution; in order to discard this factor as a cause of diversity in the reconstructed connectome, the seed areas included in the tracer connectome are the same as the ones included in the diffusion-MRI connectome. As for the diffusion-MRI connectome, the self-connections were excluded and the connection strengths were normalized between 0 and 1. The tracer connectome is shown in Figure 1D.

To evaluate the impact of introducing connections of the left SSs obtained injecting the tracing compound in the left structure (and not in the right structure as generally done in the building procedure) we built tracer-based connectome using the information of just one experiment per area (Figure S6b). In particular in Figure 4B we evaluate how reconstructing left SSs connections using different experiments (14 injection experiments performed in the right SSs and 1 injection experiment in the left SSs) impact the Predictive Power of the tracer connectome.

### 4. Surrogate connectomes

Connectomes derived with different methodologies (e.g. tracer experiments, deterministic or probabilistic diffusion-MRI tractography) give rise to different simulated resting state dynamics. Since in this study we always use the same large-scale model to simulate the functional brain patterns (reduced Wong Wang model in the bistable configuration, see section simulated dynamics), the observed differences are determined uniquely by the different structural organization used to conceptualize the brain network, i.e. the connectome.

In order to test different hypotheses about what could be the connectivity properties that give rise to the observed discrepancies in the simulated dynamics, we built different kinds of surrogate connectomes as described in what follows.

#### Averaged connectome: the role of individual variability

In order to assess the role of individual variability in dMRI data, we built an averaged connectome, both for deterministic and probabilistic tractography. We defined the

averaged connectome as a matrix whose entry 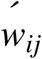, i.e. the connection strength between area *i* and area *j*, is the arithmetic mean of the values of the connection strength *w*_*ij*_ of the N individual dMRI connectomes containing both area *i* and area *j*:

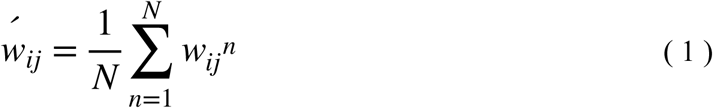

where *n* is the connectome index.

#### Filtered connectome: the role of long range connections

Comparing the connectomes in Figure 1B-D it is possible to notice that the number of long range connections detected with probabilistic, and more dramatically with deterministic, tractography is drastically lower than the one retrieved with the tracer method. It is well known that the accuracy of fiber reconstruction with diffusion-MRI data decreases with fiber distance; however, it is still unclear how to address this methodological limitation.

In order to quantify the impact of long-range connections presence in the simulated system, we filtered down the tracer connectome by removing all the connection not present in the deterministic diffusion-MRI connectomes. The filtered tracer connectome is shown in Figure 2A.

#### Symmetrized and asymmetrized connectome: the role of fiber directionality

The incapacity to detect fiber directionality is one of the main drawbacks of dMRI method.

In order to understand the influence of this property in the simulated system, we symmetrized the tracer connectome and we asymmetrized the diffusion-MRI connectome.

#### Symmetrized tracer connectome

For each asymmetric matrix exists one, and only one, decomposition that enables us to find the corresponding symmetric matrix: each generic matrix A can be decomposed in its symmetric and asymmetric part as:

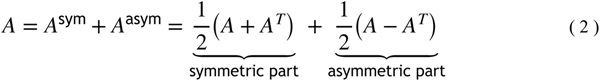

thus, symmetrizing a matrix means neglecting its asymmetric part.

Following this consideration, the tracer symmetric connectome was defined as the matrix whose entries 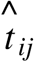 are defined as:

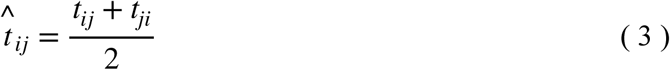

where *t*_*ij*_ represents the original tracer connection strength between area *i* and area *j*.

The symmetric tracer structural connectivity is shown in Figure 2A.

#### Asymmetrized dMRI connectome

As opposed to symmetrizing a matrix which is a straightforward procedure, asymmetrizing a matrix is an ill-posed problem, since it means introducing a new degree of freedom in the system, and not a unique solution exists. Thus, to find the asymmetric version of the dMRI connectome we assumed some constraints: we injected in each connection the same degree of asymmetry contained in the respective tracer connection, while preserving the dMRI weight balancing. In other words, our asymmetrization method assumes that the degree of asymmetry is independent on the connection strength value.

We defined the asymmetry degree *μ*_*ij*_ between connection *i* and connection *j* as:

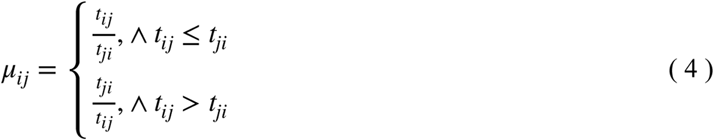

so that:

if the *ij* connection is symmetric: *t*_*ij*_ = *t*_*ji*_ ⟹*μ*_*ij*_ = + 1

if the *ij* connection is anti-symmetric: *t*_*ij*_ = − *t*_*ji*_ ⟹*μ*_*ij*_ = − 1

However, since the connection strengths in the connectome are always positively defined, *μ*_*ij*_ is a value always between 0 and 1.

The information on the directionality of the tracer connection between area *i* and area *j*, measured by *μ*_*ij*_, are inserted in the diffusion-MRI connectome by modifying the original connection *w*_*ij*_ in 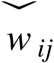:

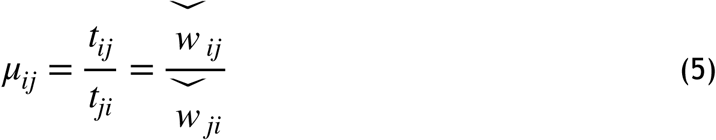

Specifically, we defined 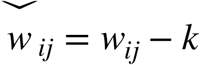 and 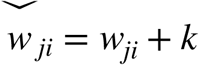, where *k* is defined as:

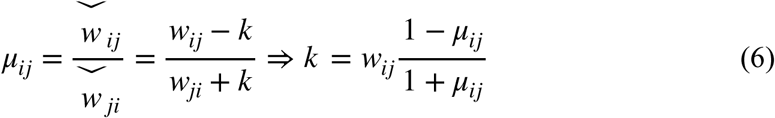

It is important to notice that the asymmetrization of the connectome does not imply the introduction of new connections: if the original diffusion-MRI connection *w*_*ij*_ is absent it follows, from the last equation, that also the increment *k* will be zero.

The asymmetrized deterministic connectome is shown in Figure 2B.

#### Hybrid connectome: the role of individual connections

We aimed to study the influence of the technique, the dMRI or the tracer one, in reconstructing the connections of a specific brain area. For this purpose, we built surrogate connectomes where all the brain wirings were reconstructed with deterministic dMRI except the connections of the region under examination that were measured with anatomical tracing.

In particular, for each deterministic dMRI connectome *W*, composed of N brain areas, we generated *N* different connectomes *W*^*k*^ by substituting the incoming and outgoing non-zero dMRI connections of area *k* with the corresponding tracer connections. The entry 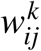 of the hybrid connectome *W*^*k*^ are defined as:

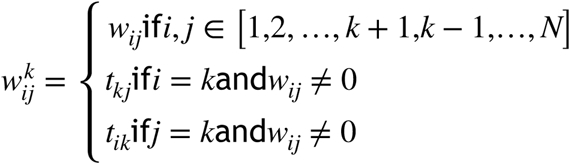

where *w*_*ij*_ and *t*_*ij*_ represent the connection strength of the original-individual deterministic dMRI and the original tracer connectome, respectively.

It is important to notice that this operation does not imply the introduction of new connections.

### 5. Comparing anatomical connectivities

#### U-static as a measure of connectome similarity

We used the Mann-Whitney test to check if the connections strength of connectomes *W*_*i*_ and *W*_*j*_ come from the same distribution. The null hypothesis of the test, *H*_0_, is that the probability of an observation, i.e. a connection strength, of the connectome *W*_*i*_ exceeding an observation from population *W*_*j*_ equals the probability of an observation *W*_*j*_ exceeding an observation from sample *W*_*i*_:

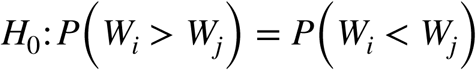

the alternative hypothesis, *H*_1_, is:

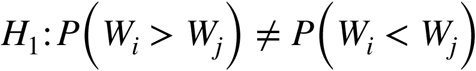

The test involves the calculation of a statistic, usually called U.

For sample size above 20, which is our case, the distribution of the U variable under the null hypothesis can be approximated using the normal distribution. The U variable ranges between 0 and *n*_1_*n*_2_, where *n*_1_ and *n*_2_ are the dimensionalities of the two connectomes. For *U* ≤ *U*\* = *n*_1_*n*_2_ /2 the test states that the *H*_0_ can be rejected.

It follows that it is possible to normalize the U value between 0 and 1, by dividing it by the product of the dimensionality of the two connectomes; in this case the discriminator value *U*\* is 0.5.

#### Euclidean distance as quantification of hemispheric functional lateralization

We quantified the functional lateralization of a given region *x* as the Euclidean distance between the functional connections of the left area *x* and the functional connections of the right area *x*.

### 6. Simulated resting state dynamics

Using the previously described connectomes we conceptualized the mouse brain as a neuronal network. The mean activity of each brain region, i.e. the network’s node, was defined by the reduced Wong Wang model (23). In this approach, the dynamics of a region is given by the whole dynamics of excitatory and inhibitory populations of leaky integrate-and-fire neurons interconnected via NMDA synapses. Here we take into account the model with a further reduction performed in (13): the dynamics of the output synaptic NMDA gating variable of the *i-*th brain area is strictly bound to the collective firing rate *H*_*i*_. The resulting model is given by the following coupled equations:

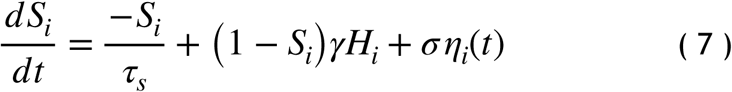

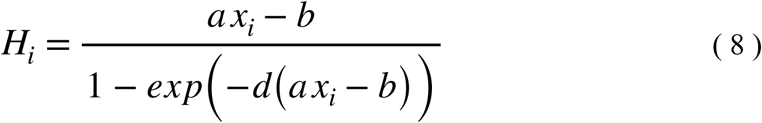

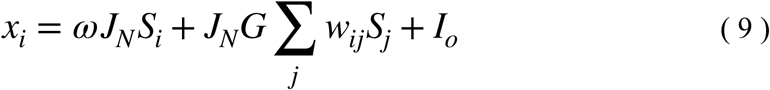

where *x*_*i*_ is the synaptic input to the *i*-th region. is a kinetic parameter fixed to 0.641, *τ*_*s*_ is the NMDA decay time constant and its value is 100 ms; *a, b* and *d* are the parameters of the input and output function *H* and are respectively equal to 270 *nC*^−1^, 108 Hz, 0.154 s. *J*_*N*_ = 0.2609*n A* is an intensity scale for the synaptic input current. *ω* is the local excitatory recurrence and *I*_*o*_ is the external input current. *G* is the coupling strength i.e. a scalar parameter which scales all the connection strengths *w*_*ij*_ without altering the global topology of the network. We set the noise amplitude σ of the normally distributed stochastic variable *η*_*i*_ to 0.015 since this level of noise is able to sustain brain states oscillations.

The external input current, *I*_*o*_, and the local excitatory recurrence, *ω*, are set to 0.3 nA and 1, respectively, in order to enrich the non-linearity of the dynamics of each brain region. In this case, studying the dynamics of isolated brain areas (*G* = 0 in equation (9)), it is possible to notice that each brain area is in a bistable state and it oscillates between high and low activity fixed points (14). It has been noticed in (14) that enriching the non-linearity of each brain areas introduces global network’s attractors that are not in trivial relation with the anatomical connectivity; this model offers the chance to reproduce the non-stationary features of the functional connectivity patterns, as shown by the checkboard pattern of the simulated FCD in Figure S2b.

For each connectome, we identified the coupling strength values for which the system is experiencing multistability. The optimal coupling strength range is defined as the values for which the system low and high states coexist, and it is identified by building the system’s bifurcation diagram as described in (13).

The brain activity, for each connectome, is simulated for 40 values of coupling strength that equally span between 0 and M, where M corresponds to the coupling strength value for which the low state (identified with the previous method), disappears. The simulations obtained from each connectome, for different coupling strength value, are used to calculate the predictive power of the connectome as explained in the section.

#### Integration scheme and BOLD signals

Model equations are numerically solved using the Euler Maruyama integration method with a fixed integration step of 0.1 ms. Simulated BOLD signal is obtained by converting the simulated synaptic activity (equation (7)) using the Balloon-Windkessel method (64) with the default value implemented in The Virtual Brain (65).

The BOLD time-series are down-sampled to 2.5 sec according to the temporal resolution of the experimental data.

### 7. Resting state signals analysis

Functional connections in the experimental and simulated time-series are explored from both spatial and temporal point of views using the Functional Connectivity (FC) and the Functional Connectivity Dynamics (FCD), respectively. We also explored the relation between functional links by estimating the Functional Meta-Connectivity (FMC).

#### Functional Connectivity (FC)

The FC matrix is defined as the matrix whose *ij*-th element is the Pearson correlation between the BOLD signal of the brain region *i* and of the brain region *j*. An example of empirical and simulated FC is shown in Figure S1.

#### Functional Connectivity Dynamics (FCD)

The FCD matrix for the experimental and simulated signals is calculated using the sliding windows approach (14, 24).

To estimate the FCD, the entire BOLD time-series is divided in time windows of a fixed length (2 min) and with a spanning of 2.5 sec; the data points within each window centered at the time *t*_*i*_were used to calculate FC(*t*_*i*_).

The *ij-*th element of the FCD matrix is calculated as the Pearson correlation between the upper triangular part of the FC(*t*_*i*_) matrix arranged as a vector and the upper triangular part of the FC(*t*_*j*_) matrix arranged as a vector. In order to observe signal correlations at frequency greater than the typical one of the BOLD signals, the sliding window length is fixed to 2 min, since, as demonstrated by (66), the non-spurious correlations in the FCD are limited by high-pass filtering of the signals with a cut-off equal to the inverse of the window length.

An example of empirical and simulated FCD is shown in Figure S2.

The typical FCD matrix during resting-state has a checkboard appearance (see experimental FCD in Figure S2) indicating that the system is switching between stable networks configuration (14, 24). We quantified the *switching degree* of the simulated and experimental system as the variance of the triangular part of the FCD once excluded the overlapping entries (i.e. the entries of the FCD matrix that quantify the correlation of FCs calculated over the sliding window of overlapping time interval). We called this quantity clue of switching (cs).

#### Functional Meta-Connectivity (FMC)

To compare the dynamical evolution of the functional connections between different systems we calculate, for each system, the FMC. The FMC, of a BOLD signals of *N* areas, is a *N* ^2^×*N* ^2^ matrix that quantifies the inter-region functional correlation of the system. The *ij*-th element of the FMC represents the Pearson correlation between the temporal evolution of the *i*-th functional link and the *j-*th functional link.

### 8. Comparing experimental and simulated BOLD signals

We quantified the ability of a given connectome to be used as a skeleton of the virtual system by comparing the accordance between the simulated functional connections, generated using that connectome, and the functional connections experimentally recorded in the resting state sessions.

As discussed before, we used the FC as the metric for quantifying the experimental and simulated functional connections. Indeed, although the FC metric is not able to capture the non-stationary nature of the resting state signals, the static functional connections are stable across resting state recordings in the same animal; on the other hand, FMC, that is able to quantify the dynamical evolution of the functional connections, is not sufficiently stable across resting state recordings (see Figure S2), and thus cannot be used for quantifying the goodness of the simulated activity.

We use 120 experimental resting state sessions recorded in 19 different animals, thus 120 experimental FCs (eFCs). For each experimental resting state session, *d*, recorded in mouse, *m*, we built a corresponding virtual mouse brain able to mimic resting state activity:

A. In the case of not-individual connectomes (i.e. the Allen SC, surrogate and original ones, and the averaged dMRI connectome) for each recording session, irrespective to the mouse scanned, we build a virtual mouse brain using the same connectome. It follows that we have 120 virtual mouse brains that are identical from the structural network point of view; however it is important to remark that virtual mouse brain with identical anatomical structure are not necessarily identical from a dynamical point of view since changing the parameters of the model one can obtain different resting state dynamics. It is this variety in dynamics that allows us to model 120 different resting state sessions using the same anatomical structure.
B. Instead, in the case of personal dMRI connectomes, surrogate and original ones, we used the information of mouse *m* to simulate the corresponding experimental resting state session. It follows that we have 120 virtual mouse brains defined according to 19 different connectomes, and thus, differently from case A, here virtual mouse brains are different both from the anatomical and dynamical point of view.

In these virtual brains, we simulate resting state activity using the reduced Wong Wang model in the bistable configuration. In the model there is a parameter, i.e. the coupling strength G (eq. 9), whose value is not fixed by biological constraints, but is variable and is used to optimize the simulated output (12, 14, 55). For each recording session, thus for each eFC, we optimize the coupling strength value in order to maximize the correlation between the eFC and the simulated FC (sFC). In particular, for each connectome, we explored 40 different values of the coupling strength in an equally spanned interval range. The interval range is identified according to the bifurcation diagram of each connectome built as described in (13).

For each mouse, *m*, and each session, *d*, we defined the *PP* of a given connectome *c* as the maximum Pearson correlation between the empirical FC (eFC) and the simulated FC (sFC) obtained for the different coupling strength values G:

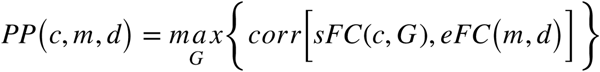

The PP of a given connectome *PP*(*c*) is the mean over all the mice and the sessions of the *PP*(*c, m, d*):

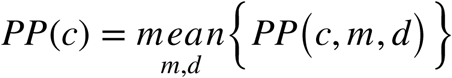

Since 120 experimental resting state sessions enter in the analysis, it follows that for each type of connectome *c* the average PP is calculated over 120 PP values. In order to assess the significance of the difference in PP of differently derived connectomes we used the p-value calculated through the Welch’s test; we corrected the p-values for multiple comparisons using the Bonferroni correction. We measure the effect size using the Cohen’s d and we calculated the 95% confidence intervals (CIs) using the estimation stats framework as described in (67) and available at https://www.estimationstats.com/.

Finally, we want to point out that the simulated functional network is composed of more areas than the experimental one since the simulation is based on the anatomical information that has a greater spatial resolution than the functional one. Thus, in order to correlate the eFC and the sFC we reduced them to the same number of areas.

### 9. Data sharing

All the data and the codes are available on request.

---

**Table S1:**
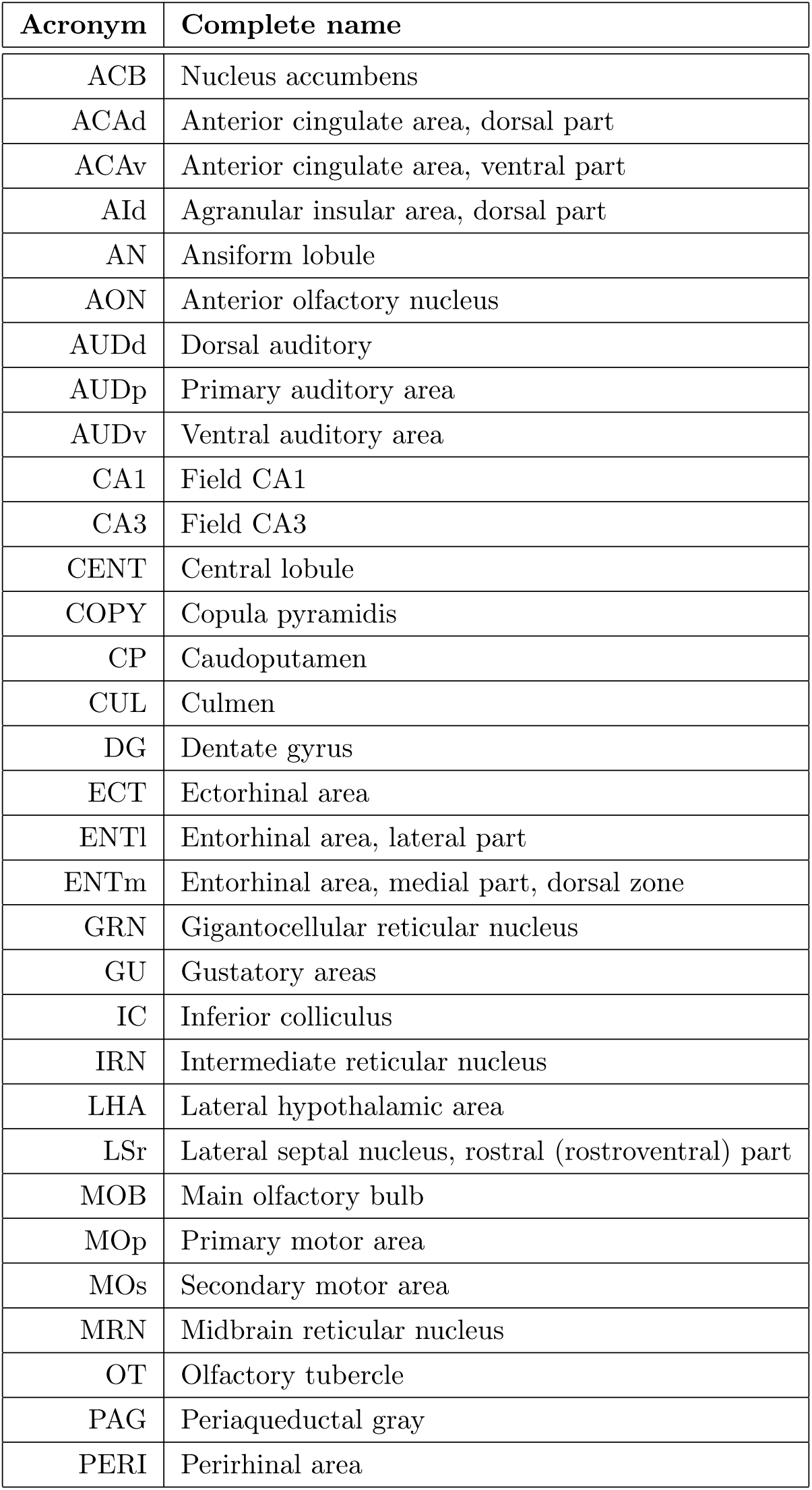
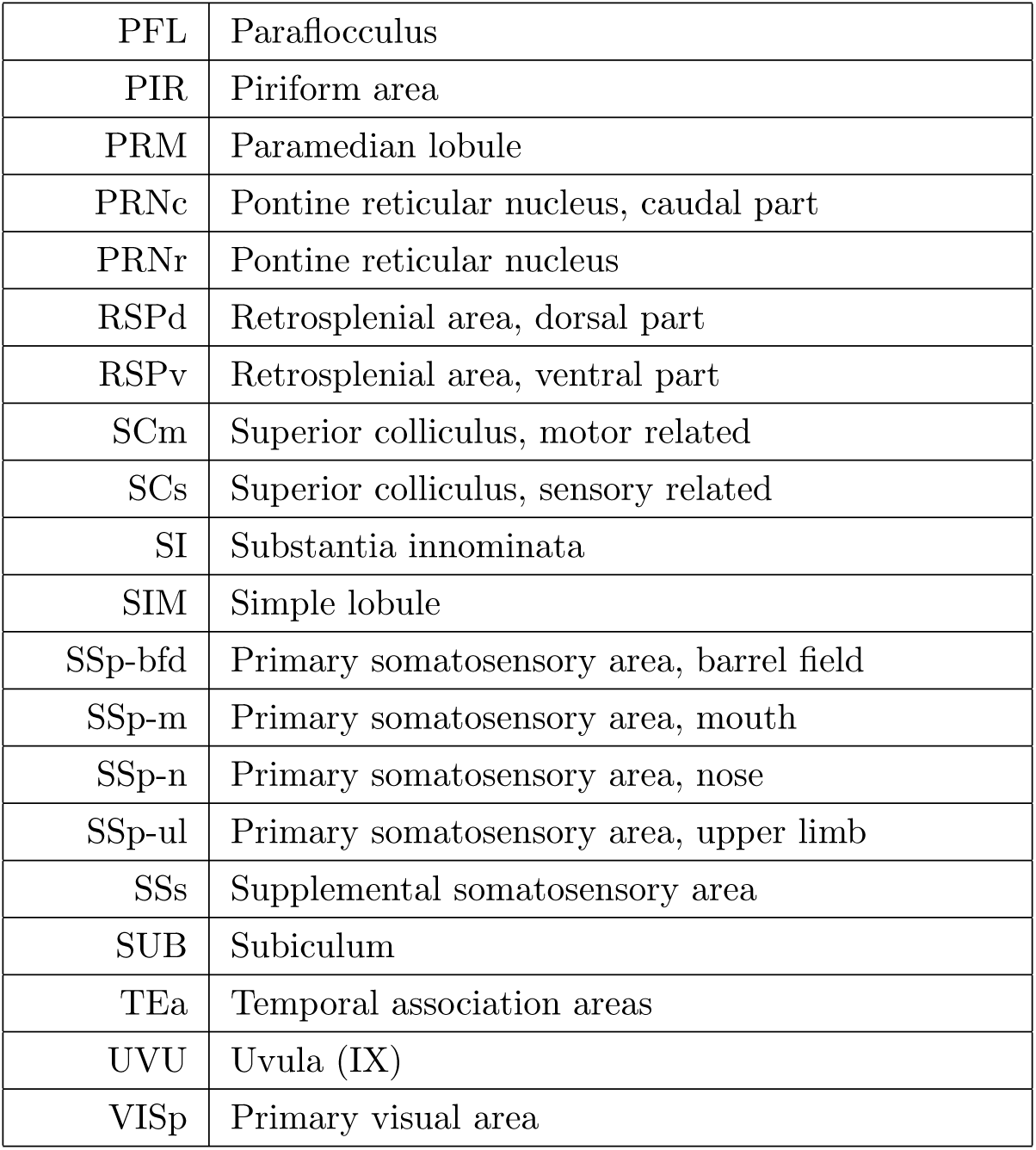
List of the acronyms for the different brain regions considered in this work.

| Acronym | Complete name |
| --- | --- |
| ACB | Nucleus accumbens |
| ACAd | Anterior cingulate area, dorsal part |
| ACAv | Anterior cingulate area, ventral part |
| AId | Agranular insular area, dorsal part |
| AN | Ansiform lobule |
| AON | Anterior olfactory nucleus |
| AUDd | Dorsal auditory |
| AUDp | Primary auditory area |
| AUDv | Ventral auditory area |
| CA1 | Field CA1 |
| CA3 | Field CA3 |
| CENT | Central lobule |
| COPY | Copula pyramidis |
| CP | Caudoputamen |
| CUL | Culmen |
| DG | Dentate gyrus |
| ECT | Ectorhinal area |
| ENTl | Entorhinal area, lateral part |
| ENTm | Entorhinal area, medial part, dorsal zone |
| GRN | Gigantocellular reticular nucleus |
| GU | Gustatory areas |
| IC | Inferior colliculus |
| IRN | Intermediate reticular nucleus |
| LHA | Lateral hypothalamic area |
| LSr | Lateral septal nucleus, rostral (rostroventral) part |
| MOB | Main olfactory bulb |
| MOp | Primary motor area |
| MOs | Secondary motor area |
| MRN | Midbrain reticular nucleus |
| OT | Olfactory tubercle |
| PAG | Periaqueductal gray |
| PERI | Perirhinal area |
| PFL | Paraflocculus |
| PIR | Piriform area |
| PRM | Paramedian lobule |
| PRNc | Pontine reticular nucleus, caudal part |
| PRNr | Pontine reticular nucleus |
| RSPd | Retrosplenial area, dorsal part |
| RSPv | Retrosplenial area, ventral part |
| SCm | Superior colliculus, motor related |
| SCs | Superior colliculus, sensory related |
| SI | Substantia innominata |
| SIM | Simple lobule |
| SSp-bfd | Primary somatosensory area, barrel field |
| SSp-m | Primary somatosensory area, mouth |
| SSp-n | Primary somatosensory area, nose |
| SSp-ul | Primary somatosensory area, upper limb |
| SSs | Supplemental somatosensory area |
| SUB | Subiculum |
| TEa | Temporal association areas |
| UVU | Uvula (IX) |
| VISp | Primary visual area |

**Figure S1:**
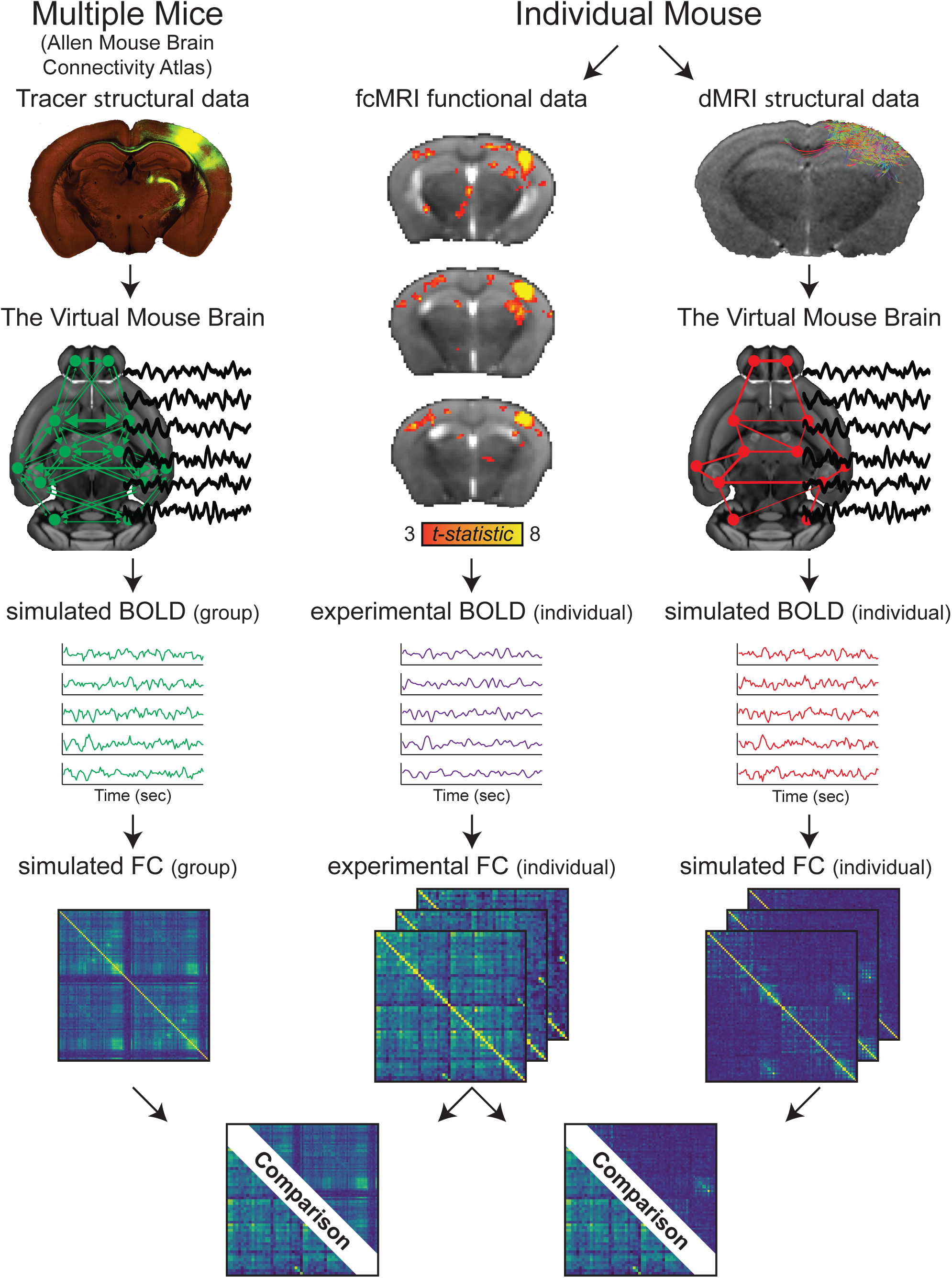
Experimental workflow. Each mouse is scanned to obtain both SC and FC data. First, FC is estimated empirically for each subject as illustrated by the seed-based analysis of the barrel-related primary somatosensory cortex (t-test of 7 recording sessions of the same mouse, p < 0.01, uncorrected, voxel extent = 20) and FC matrices. Then, simulated BOLD activity is generated by the virtual mouse brain using dMRI-based SC. The simulated and experimental brain dynamics are compared through a static-FC metric to estimate the predictive power. Then, the results are compared to the predictive power of the gold standard Allen SC. Note that the Allen SC, but not dMRI-based SC, provides fiber directionality.

**Figure S2:**
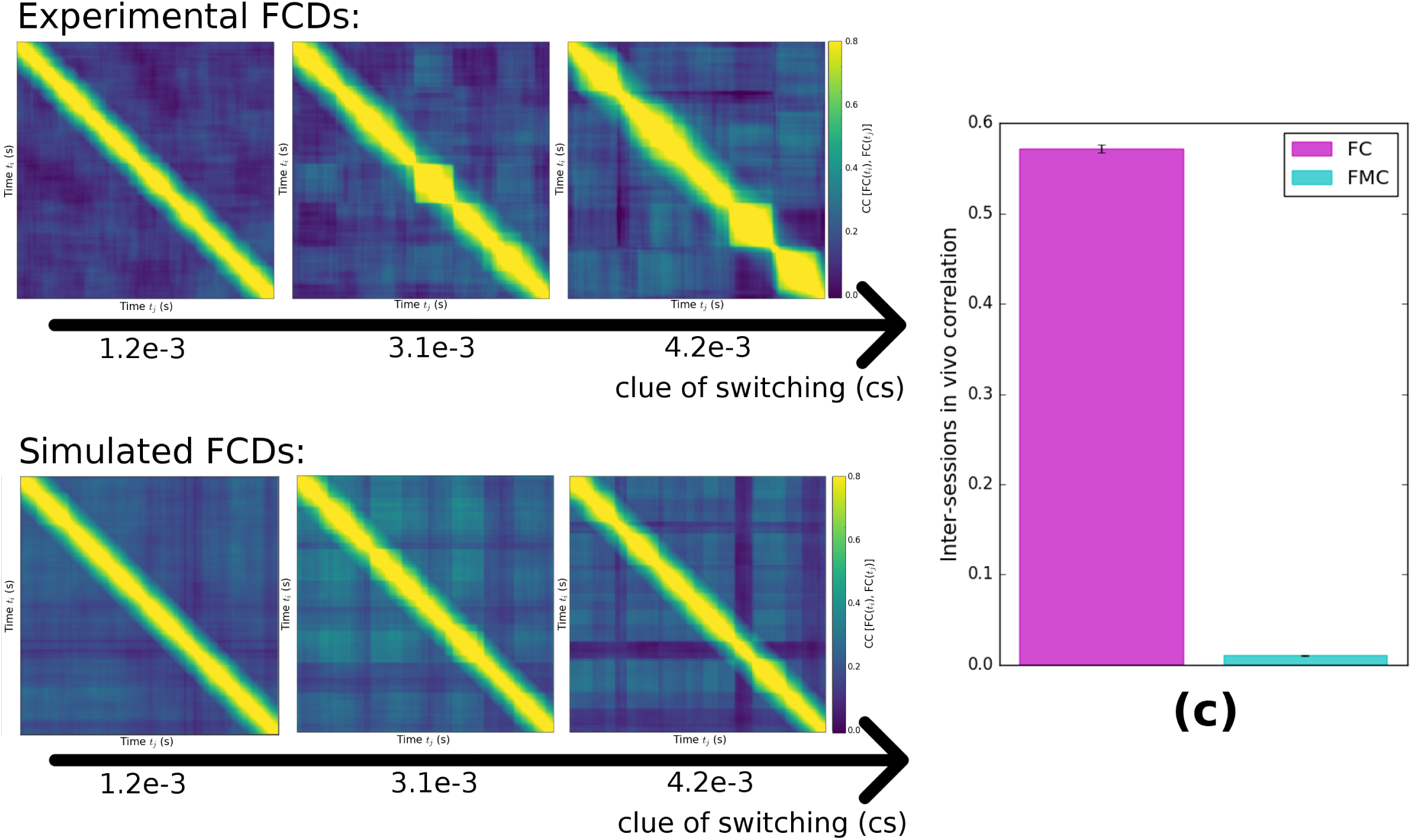
Variability of FCD metric in experimental (a) and simulated (b) data. FCDs on the top are calculated from the experimental resting state data from 3 different scanning sessions; FCDs on the bottom are calculated from simulated resting state data; in both case we use a sliding window length of 2 minutes and a spanning of 2.5 seconds. We quantify the presence of the switching, i.e. the checkboard pattern in the FCD matrix, as the variance of the triangular part of the FCD, once excluded the overlapping entries. We call this quantity: clue of switching (cs). The cs value is indicated below each FCD. FCDs are ordered for increasing cs values. The checkboard pattern appears more clearly as cs increases (c) The height of the bar represents the Pearson correlation between inter-sessions for experimental FC (magenta bar) and FMC (blue bar). The result shows that the FMC matrix can not be considered as a metric since FMC is poorly reproducible across sessions in the same animal.

**Figure S3:**
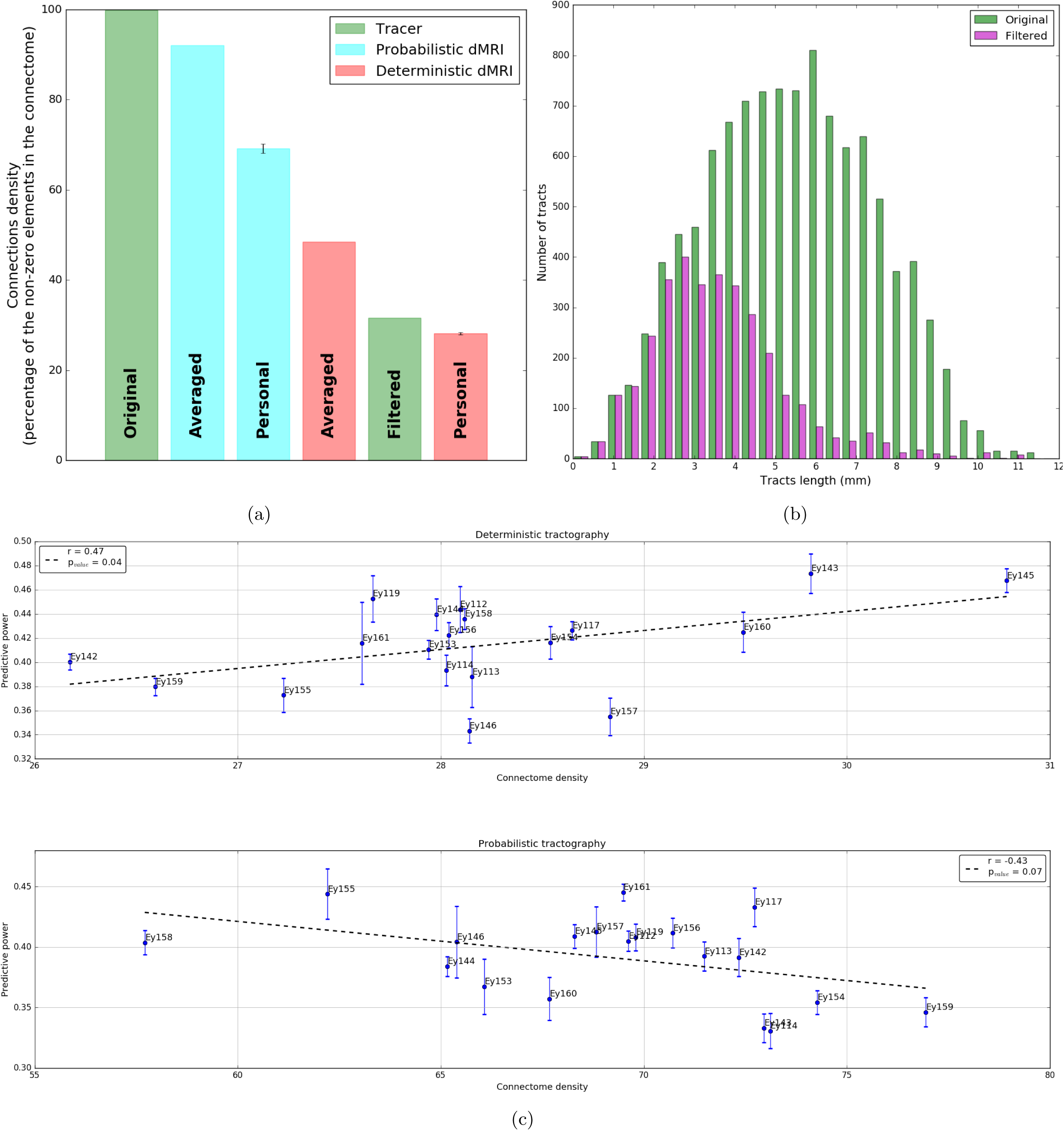
(a) The height of the bars represents the density of the connectome used in this study. The difference in the rank of the bars in this plot and those representing the connectomes PP (Figure 2-3) highlights the absence of a relation between the connectome’s density and its PP. In line with this observation, the plots in (c) show the relation between the PP and the density of the dMRI-based connectomes processed with the deterministic tractography (plot on the top) and probabilistic tractography (plot in the bottom).The relation between the connectomes density and the PP it is opposite between the case of deterministic and probabilistic processing. In the case of deterministic processed tractography data, denser connectomes, i.e. with less false negative, have a greater PP than sparser connectomes, that is connectomes with more false negative. Conversely, in the case of probabilistic processed data, sparser connectomes, i.e. with less false negative, have a greater PP than denser connectome. From the results of the panels (a) and (c) it follows that the number of connections in a connectome does not directly relate with its ability in predicting brain dynamics. The histogram in (b) shows the distribution of the fibers lengths included in the original tracer-based (green bars) and in the filtered tracer-based (magenta bars) connectome. The filtered connectome was obtained from the original tracer connectome by removing the connections not present in at least one of the 19 deterministic connectomes (68% of the tracer-based connections removed). This operation results in removing mainly long-range connections (the mean length of the tracts contained in the original tracer connectome and in the filtered one is respectively 5.40*±*0.02 mm and 3.57*±*0.03 mm).

**Figure S4:**
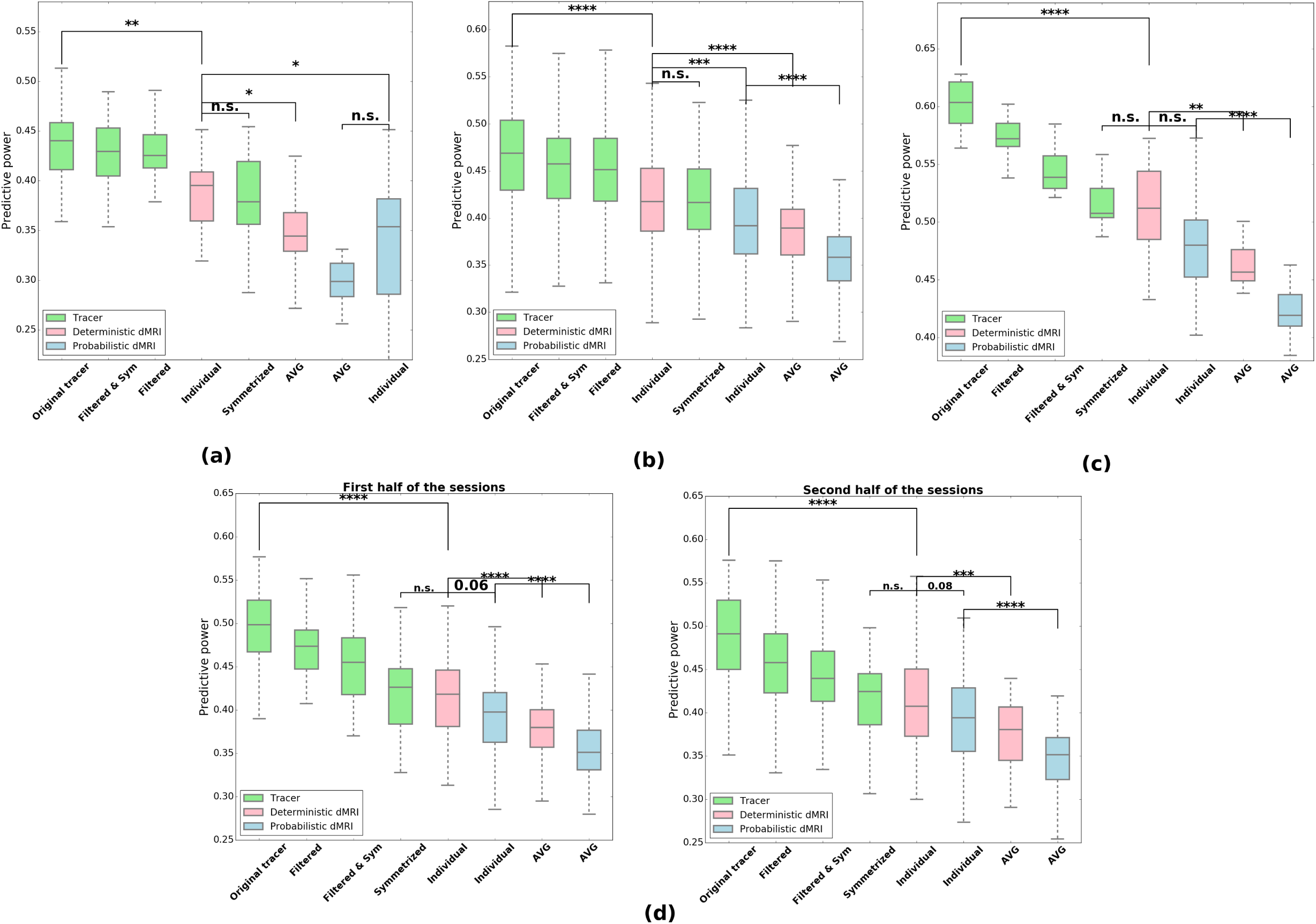
The box plots represent the predictive power of the tracer-based (green bars), deterministic dMRI-based (red bars) and probabilistic dMRI-based (blue bars) connectomes. The meaning and the construction of the figures are analogous to Figure 3b 3C in the main text. (a) Results obtained for examination of 7 wild type inbred C57BL/6 mice (4*±*2 sessions per animal, 11*±*4 minutes per session, mean*±*SD). The results are similar to those found for in hybrid mice. (b) Results for hybrid mice obtained using experimental resting state data preprocessed using global signal regression. The results are consistent to those presented in Figure 2 and Figure 3, that are those obtained from experimental resting state data not not globally signals regressed. (c) Results obtained after averaging recording sessions in the same animal. The predictive power trend is analogous to the one obtained considering separately the contribution of each recording session. (d) The two box plots show the predictive power calculated splitting the recoding sessions in two; results are consistent with the one obtained considering the complete dataset.

**Figure S5:**
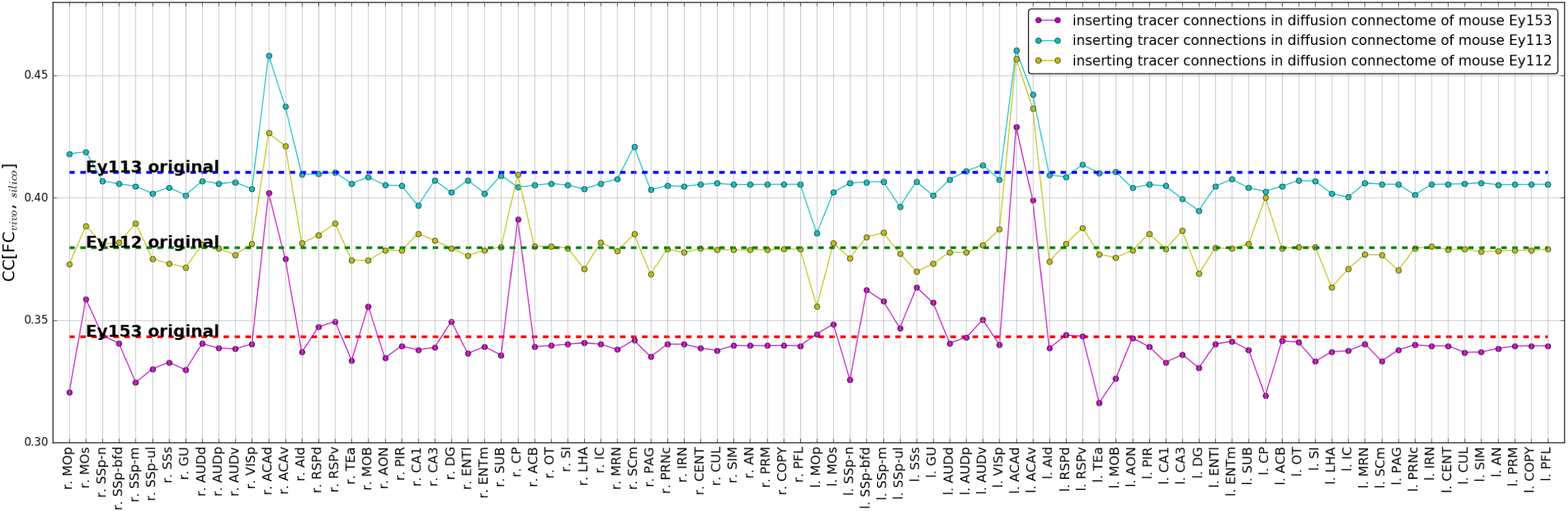
Dots of different colors represent the predictive power of the deterministic dMRI connectome of different mice (namely Ey113, Ey112 and Ey153) with the connections of the brain area (x-axis) replaced with the corresponding tracer connections. Red, green and blue dashed lines represent the predictive power of the personal deterministic dMRI connectome of different mice. The figure shows that the change in predictive power strictly depends on the considered brain areas and on the mouse connectome: for example, the replacement of the right caudoputamen connections strongly enhances the predictive power of the deterministic dMRI connectome of mice Ey153 and Ey112, but not of mouse Ey113. Conversely, the replacement of cerebellum’s connections does not really affect the performance of the connectome in predicting brain dynamics. The labels of brain areas are shown in table S1.

**Figure S6:**
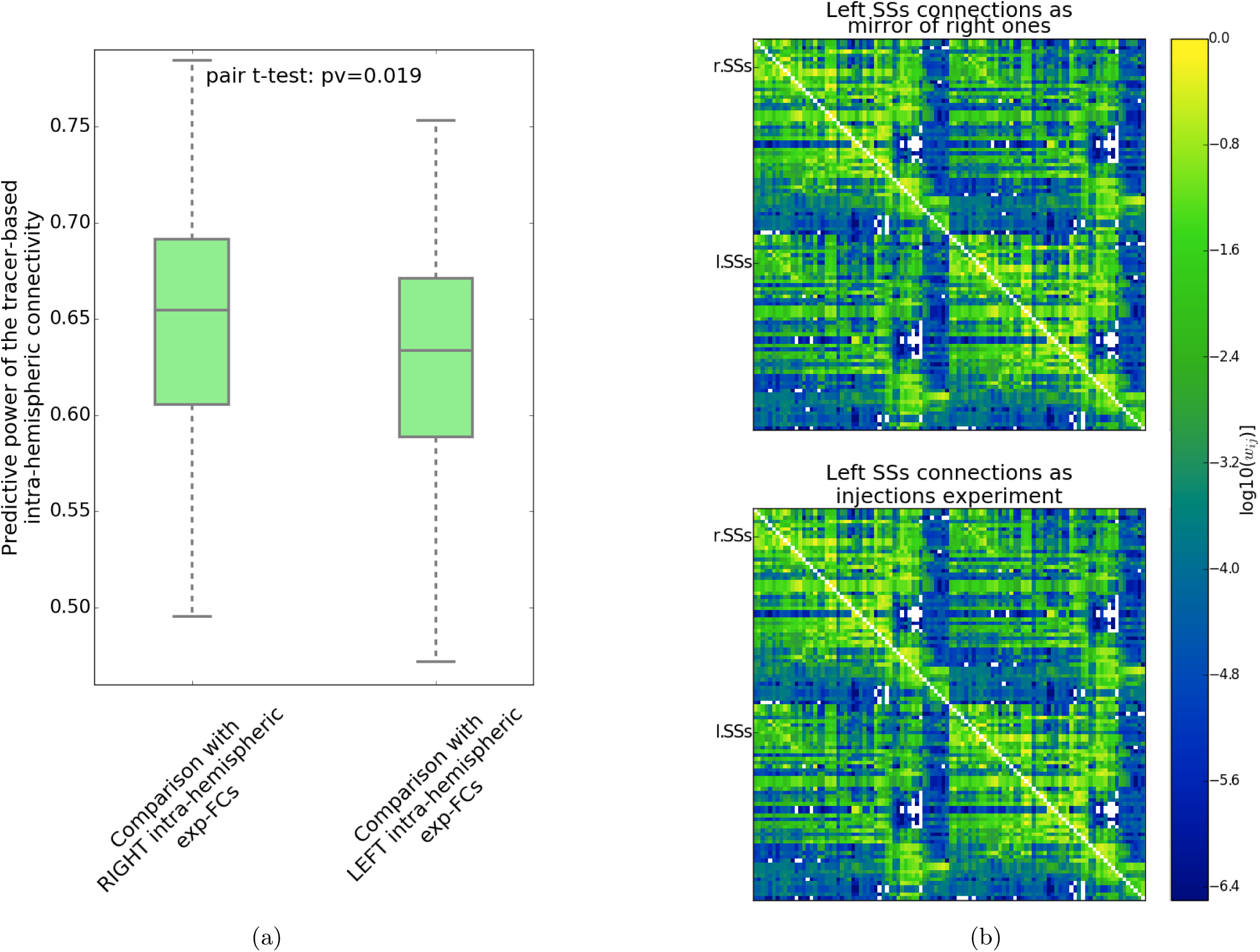
(a)The box plot represent the predictive power of the tracer-based connectome built using only intra-hemispheric connections. The predictive power is estimated comparing the predictions of the connectome with the intra-hemispheric experimental functional connections measured from the right (left bar) or the left (right bar) hemisphere. The difference in the predictive power shows that intra-hemispheric tracer-based connections, obtained after injecting the compound in the right hemisphere, are able to predict better right than left intra-hemispheric functional connections. The result suggests that the mouse brain is lateralized. (b) Tracer-based connectomes built using the information of one injection experiment per area. The difference between the two connectomes relies on the definition of the connections of the left SSs: in the connectome on the top, the connections of the left SSs are the mirror image of the connections of the right SSs, that are connections built using information from an injection experiment performed in the right SSs region. In contrast, in the connectome shown on the bottom, the left SSs connections are built using the information of an injection experiment performed in the left SSs area.

## REFERENCES

1. Sporns O, Tononi G, Kötter R (2005) The human connectome: a structural description of the human brain. PLoS Comput Biol 1(4):e42.

2. Yeh F-C, et al. (2016) Quantifying differences and similarities in whole-brain white matter architecture using local connectome fingerprints. PLoS Comput Biol 12(11):e1005203.

3. Mikhael SS, et al. (2019) Manually-parcellated gyral data accounting for all known anatomical variability. Sci Data 6:190001.

4. Biswal B, Zerrin Yetkin F, Haughton VM, Hyde JS (1995) Functional connectivity in the motor cortex of resting human brain using echo-planar mri. Magn Reson Med 34(4):537–541.

5. Finn ES, et al. (2015) Functional connectome fingerprinting: identifying individuals using patterns of brain connectivity. Nat Neurosci 18(11):1664–1671.

6. Gratton C, et al. (2018) Functional brain networks are dominated by stable group and individual factors, not cognitive or daily variation. Neuron 98(2):439–452.

7. Mueller S, et al. (2013) Individual variability in functional connectivity architecture of the human brain. Neuron 77(3):586–595.

8. Finn ES, Todd Constable R (2016) Individual variation in functional brain connectivity: implications for personalized approaches to psychiatric disease. Dialogues Clin Neurosci 18(3): 277–287.

9. Sanz Leon P, et al. (2013) The Virtual Brain: a simulator of primate brain network dynamics. Front Neuroinformatics 7:10.

10. Ghosh A, Rho Y, McIntosh AR, Kötter R, Jirsa VK (2008) Noise during rest enables the exploration of the brain’s dynamic repertoire. PLoS Comput Biol 4(10):e1000196.

11. Jirsa VK, Jantzen KJ, Fuchs A, Kelso JAS (2002) Spatiotemporal forward solution of the EEG and MEG using network modeling. IEEE Trans Med Imaging 21(5):493–504.

12. Deco G, Jirsa VK, McIntosh AR (2011) Emerging concepts for the dynamical organization of resting-state activity in the brain. Nat Rev Neurosci 12(1):43.

13. Deco G, et al. (2013) Resting-state functional connectivity emerges from structurally and dynamically shaped slow linear fluctuations. J Neurosci 33(27):11239–11252.

14. Hansen EC, Battaglia D, Spiegler A, Deco G, Jirsa VK (2015) Functional connectivity dynamics: modeling the switching behavior of the resting state. Neuroimage 105:525–535.

15. Jirsa VK, et al. (2017) The Virtual Epileptic Patient: Individualized whole-brain models of epilepsy spread. NeuroImage 145:377–388.

16. Wedeen VJ, et al. (2008) Diffusion spectrum magnetic resonance imaging (DSI) tractography of crossing fibers. Neuroimage 41(4):1267–1277.

17. Kötter R (2007) Anatomical Concepts of Brain Connectivity. Handbook of Brain Connectivity, Understanding Complex Systems. (Springer, Berlin, Heidelberg), pp 149–167.

18. Oh SW, et al. (2014) A mesoscale connectome of the mouse brain. Nature 508(7495):207.

19. Gămănut R, et al. (2018) The Mouse Cortical Connectome, Characterized by an Ultra-Dense Cortical Graph, Maintains Specificity by Distinct Connectivity Profiles. Neuron 97(3): 698-715.e10.

20. Ypma RJF, Bullmore ET (2016) Statistical Analysis of Tract-Tracing Experiments Demonstrates a Dense, Complex Cortical Network in the Mouse. PLoS Comput Biol 12(9):e1005104.

21. Melozzi F, Woodman MM, Jirsa VK, Bernard C (2017) The Virtual Mouse Brain: A Computational Neuroinformatics Platform To Study Whole Mouse Brain Dynamics. eNeuro:ENEURO–0111.

22. Bergmann E, Zur G, Bershadsky G, Kahn I (2016) The organization of mouse and human cortico-hippocampal networks estimated by intrinsic functional connectivity. Cereb Cortex:1–16.

23. Wong K-F, Wang X-J (2006) A recurrent network mechanism of time integration in perceptual decisions. J Neurosci 26(4):1314–1328.

24. Allen EA, et al. (2014) Tracking whole-brain connectivity dynamics in the resting state. Cereb Cortex 24(3):663–676.

25. Grandjean J, et al. (2017) Dynamic reorganization of intrinsic functional networks in the mouse brain. NeuroImage 152:497–508.

26. Laumann TO, et al. (2015) Functional system and areal organization of a highly sampled individual human brain. Neuron 87(3):657–670.

27. Van Dijk KRA, et al. (2009) Intrinsic Functional Connectivity As a Tool For Human Connectomics: Theory, Properties, and Optimization. J Neurophysiol 103(1):297–321.

28. Tournier JD, Calamante F, Connelly A (2010) Improved probabilistic streamlines tractography by 2nd order integration over fibre orientation distributions. Proceedings of the International Society for Magnetic Resonance in Medicine, p 1670.

29. Tournier J-D, Calamante F, Connelly A (2012) MRtrix: diffusion tractography in crossing fiber regions. Int J Imaging Syst Technol 22(1):53–66.

30. Zalesky A, et al. (2016) Connectome sensitivity or specificity: which is more important? Neuroimage 142:407–420.

31. Fox MD, Zhang D, Snyder AZ, Raichle ME (2009) The global signal and observed anticorrelated resting state brain networks. J Neurophysiol 101(6):3270–3283.

32. Craddock RC, et al. (2013) Imaging human connectomes at the macroscale. Nat Methods 10(6): 524–539.

33. Li L, Rilling JK, Preuss TM, Glasser MF, Hu X (2012) The effects of connection reconstruction method on the interregional connectivity of brain networks via diffusion tractography. Hum Brain Mapp 33(8):1894–1913.

34. Kale P, Zalesky A, Gollo LL (2018) Estimating the impact of structural directionality: How reliable are undirected connectomes? Netw Neurosci 2(02):259–284.

35. Mohajerani MH, et al. (2013) Spontaneous cortical activity alternates between motifs defined by regional axonal projections. Nat Neurosci 16(10):1426.

36. Stafford JM, et al. (2014) Large-scale topology and the default mode network in the mouse connectome. Proc Natl Acad Sci 111(52):18745–18750.

37. Díaz-Parra A, Osborn Z, Canals S, Moratal D, Sporns O (2017) Structural and functional, empirical and modeled connectivity in the cerebral cortex of the rat. Neuroimage 159:170–184.

38. Zimmermann J, Griffiths J, Schirner M, Ritter P, McIntosh AR (2018) Subject specificity of the correlation between large-scale structural and functional connectivity. Netw Neurosci 3(1):90–106.

39. Farde L, Plavén-Sigray P, Borg J, Cervenka S (2018) Brain neuroreceptor density and personality traits: towards dimensional biomarkers for psychiatric disorders. Philos Trans R Soc B Biol Sci 373(1744):20170156.

40. Rangaprakash D, Wu G-R, Marinazzo D, Hu X, Deshpande G (2018) Hemodynamic response function (HRF) variability confounds resting-state fMRI functional connectivity. Magn Reson Med 80(4):1697–1713.

41. Desai M, et al. (2010) Mapping brain networks in awake mice using combined optical neural control and fMRI. J Neurophysiol 105(3):1393–1405.

42. Schlegel F, Schroeter A, Rudin M (2015) The hemodynamic response to somatosensory stimulation in mice depends on the anesthetic used: implications on analysis of mouse fMRI data. Neuroimage 116:40–49.

43. Uludağ K, Müller-Bierl B, Uğurbil K (2009) An integrative model for neuronal activity-induced signal changes for gradient and spin echo functional imaging. Neuroimage 48(1):150–165.

44. Calabrese E, Badea A, Cofer G, Qi Y, Johnson GA (2015) A diffusion MRI tractography connectome of the mouse brain and comparison with neuronal tracer data. Cereb Cortex 25(11): 4628–4637.

45. Mukherjee P, Chung SW, Berman JI, Hess CP, Henry RG (2008) Diffusion tensor MR imaging and fiber tractography: technical considerations. Am J Neuroradiol 29(5):843–852.

46. Smith RE, Tournier J-D, Calamante F, Connelly A (2012) Anatomically-constrained tractography: improved diffusion MRI streamlines tractography through effective use of anatomical information. Neuroimage 62(3):1924–1938.

47. Fritz FJ, et al. (2019) Ultra-high resolution and multi-shell diffusion MRI of intact ex vivo human brains using kT-dSTEAM at 9.4T. NeuroImage 202:116087.

48. Stephan KE, et al. (2001) Advanced database methodology for the Collation of Connectivity data on the Macaque brain (CoCoMac). Philos Trans R Soc Lond B Biol Sci 356(1412):1159–1186.

49. Markov NT, et al. (2014) A Weighted and Directed Interareal Connectivity Matrix for Macaque Cerebral Cortex. Cereb Cortex 24(1):17–36.

50. Silva AJ, et al. (1997) Mutant mice and neuroscience: recommendations concerning genetic background. Neuron 19(4):755–759.

51. Sittig LJ, et al. (2016) Genetic background limits generalizability of genotype-phenotype relationships. Neuron 91(6):1253–1259.

52. Shinohara Y, Hosoya A, Hirase H (2013) Experience enhances gamma oscillations and interhemispheric asymmetry in the hippocampus. Nat Commun 4:1652.

53. Deco G, Kringelbach ML, Jirsa VK, Ritter P (2017) The dynamics of resting fluctuations in the brain: metastability and its dynamical cortical core. Sci Rep 7(1):3095.

54. Messé A, Rudrauf D, Benali H, Marrelec G (2014) Relating structure and function in the human brain: relative contributions of anatomy, stationary dynamics, and non-stationarities. PLoS Comput Biol 10(3):e1003530.

55. Deco G, Jirsa VK, McIntosh AR (2013) Resting brains never rest: computational insights into potential cognitive architectures. Trends Neurosci 36(5):268–274.

56. Saggio ML, Ritter P, Jirsa VK (2016) Analytical operations relate structural and functional connectivity in the brain. PloS One 11(8):e0157292.

57. Fornito A, Zalesky A, Breakspear M (2015) The connectomics of brain disorders. Nat Rev Neurosci 16(3):159.

58. Calhoun VD, Lawrie SM, Mourao-Miranda J, Stephan KE (2017) Prediction of Individual Differences from Neuroimaging Data. NeuroImage 145:135–136.

59. Deco G, Kringelbach ML (2014) Great Expectations: Using Whole-Brain Computational Connectomics for Understanding Neuropsychiatric Disorders. Neuron 84(5):892–905.

60. Besson P, et al. (2017) Anatomic consistencies across epilepsies: a stereotactic-EEG informed high-resolution structural connectivity study. Brain 140(10):2639–2652.

61. Greve DN, Fischl B (2009) Accurate and robust brain image alignment using boundary-based registration. Neuroimage 48(1):63–72.

62. Tournier J-D, Calamante F, Connelly A (2007) Robust determination of the fibre orientation distribution in diffusion MRI: non-negativity constrained super-resolved spherical deconvolution. Neuroimage 35(4):1459–1472.

63. Smith RE, Tournier J-D, Calamante F, Connelly A (2013) SIFT: spherical-deconvolution informed filtering of tractograms. Neuroimage 67:298–312.

64. Friston KJ, Mechelli A, Turner R, Price CJ (2000) Nonlinear responses in fMRI: the Balloon model, Volterra kernels, and other hemodynamics. NeuroImage 12(4):466–477.

65. Sanz-Leon P, Knock SA, Spiegler A, Jirsa VK (2015) Mathematical framework for large-scale brain network modeling in The Virtual Brain. Neuroimage 111:385–430.

66. Leonardi N, Van De Ville D (2015) On spurious and real fluctuations of dynamic functional connectivity during rest. Neuroimage 104:430–436.

67. Ho J, Tumkaya T, Aryal S, Choi H, Claridge-Chang A (2019) Moving beyond P values: data analysis with estimation graphics. Nat Methods:1.

